# LIFE-seq: longitudinal spatial transcriptomics of living tissue

**DOI:** 10.64898/2026.11.13.766339

**Authors:** Wenjia Wang, Handong Wang, Kaiqiang Ye, Feiyu Ding, Liuxi Chu, Xiaoying Ma, Kaitong Dang, Liyong He, Yan Huang, Huiying Li, Zhenyu Yin, Xiangwei Zhao

**Affiliations:** State Key Laboratory of Digital Medical Engineering, School of Biological Science & Medical Engineering, Southeast University, Nanjing, 210096, China; Department of Neurosurgery, Nanjing BenQ Medical Center, The Affiliated BenQ Hospital of Nanjing Medical University, Nanjing 210019, Jiangsu Province, China; The First Affiliated Hospital of Wenzhou Medical University, Wenzhou, Zhejiang 325000, China; Department of Geriatric Oncology, Affiliated Nanjing Drum Tower Hospital of Nanjing University Medical School, Nanjing 210000, China

**Keywords:** live tissue, nanoforest, RNA sequencing (RNA-seq), temporal, spatial transcriptomics

## Abstract

Spatial transcriptomics (ST) has deepened our understanding of cellular spatial heterogeneity, yet existing technologies are predominantly confined to fixed or frozen specimens, limiting longitudinal molecular profiling of the same living tissue. To overcome this limitation, we developed LIFE-seq (Living-tissue in situ nanoforest extraction for spatiotemporal transcriptomic mapping), a minimally invasive approach that combines high-density nanoforest arrays for molecular sampling with directed transfer onto spatially barcoded messenger RNA (mRNA) capture chips. LIFE-seq enables spatial transcriptomic profiling while preserving overall tissue integrity and provides a modular strategy with the potential to integrate different 3’-end capture platforms. Performance validation demonstrated high concordance in global transcriptomic profiles and spatial gene expression patterns between nanoforest-derived replicas and conventional tissue sections, as well as faithful recapitulation of pathological features in a traumatic brain injury (TBI) model. Furthermore, longitudinal spatial transcriptomic profiling of cortical injury in the same mouse revealed temporally distinct injury responses, progressive activation of immune-associated gene programs, and spatially heterogeneous cell-type-specific expression changes. By enabling repeated molecular sampling and spatial transcriptomic analysis of living tissue, LIFE-seq provides a versatile framework for investigating tissue dynamics and pathological progression *in vivo*.

## Introduction

Spatial transcriptomics (ST) maps gene expression while preserving tissue architecture, providing important insights into complex biological systems ^1–3^. However, current mainstream ST methods are restricted to fresh-frozen and formalin-fixed, paraffin-embedded (FFPE) specimens. This precludes subsequent molecular or phenotypic assays in the same cells or tissue and prevents transcriptomic analysis of living tissue. Furthermore, fixed specimens capture cellular states at a single time point, and the observed expression profiles may simply reflect expression heterogeneity. These methods constrain observations to active transcriptional states at one time point, while other cells with similar functions may be quiescent. Their ability to infer cellular states accurately at the level of individual genes and across time therefore remains debated.

To address the challenges of live-cell transcriptomics, researchers have explored several in vitro labeling and sampling strategies. Methods based on photoactivation or region-of-interest (ROI) labeling, including transcriptome in vivo analysis (TIVA) ^4^, ZipSeq ^5^ and spatially photoactivatable color encoded cell address tags (SPACECAT) ^6^, enable spatial labeling of living cells within intact tissue. However, these approaches generally have low spatial resolution or very low analytical throughput and require demanding experimental conditions, limiting large-scale cellular mapping in complex tissues. Enzyme-mediated or photocatalytic proximity labeling methods, such as APEX-seq ^7^ and μMap-seq ^8^, preserve in situ molecular information but require genetic transfection to introduce exogenous probes. This restricts their application to a small number of readily transfected cell lines and inevitably perturbs native cellular physiology. Another emerging approach uses atomic force microscopy (AFM) ^9^ or nanotubes ^10^ to extract intracellular material directly from single cells. In this area, Live-seq was the first to enable sequential monitoring of transcriptome-wide gene expression in living cells ^11^. Nevertheless, these single-cell nanobiopsy methods generally do not provide the three-dimensional positions of cells within the tissue microenvironment, are difficult to apply in vivo, and face substantial constraints in sampling throughput and repeated sampling. Thus, current live-cell transcriptomic methods struggle to combine preservation of spatial context, maintenance of native physiology without damage, and high-throughput in vivo mapping.

Three-dimensional micro- and nanostructures with high aspect ratios offer a promising way to overcome these physical and biochemical barriers ^12^. Building on microneedle arrays, these structures combine versatility with minimal invasiveness and have advanced rapidly in drug delivery, biomarker detection and tissue repair ^13, 14^. Recently, acupuncture-extracted in vivo spatiotemporal transcriptomics (aivST) was developed by functionalizing clinical gold acupuncture microneedle arrays with spatial barcodes, enabling in vivo spatiotemporal transcriptomic tracking of skin wound healing in mice ^15^. However, direct encoding on millimeter-scale microneedles is cumbersome, spatial resolution is constrained by the physical needle pitch, and deep penetration can cause tissue damage and superposition of signals along the depth axis. Smaller nanoneedle arrays provide denser molecular capture interfaces at higher resolution with minimal cellular perturbation. For example, Spectrum-FISH, which uses silicon nanoneedle arrays, and the endoscope-integrated ENDO-Genome system have enabled bloodless in situ RNA extraction and spatial analysis at single-cell resolution in fresh tissue and human internal organs ^16, 17^. These sequencing-free strategies rely on hybridization with predefined spectrally encoded probes and target only tens of known genes per measurement, limiting unbiased transcriptome-wide exploration. Porous silicon nanoneedles can also generate molecular replicas of living brain tissue by non-destructive in situ imprinting for longitudinal mass spectrometry imaging ^18^, but this approach is restricted to lipidomics. Combining minimally invasive, non-destructive sampling, high spatial resolution and unbiased whole-transcriptome sequencing in living tissue using high-density nanoforest arrays therefore remains a key challenge.

Here we introduce LIFE-seq (Living-tissue in situ nanoforest extraction for spatiotemporal transcriptomic mapping), a method for minimally invasive in situ sampling and spatiotemporal transcriptomic analysis of living tissue. Nanoforest arrays enable minimally invasive and repeated in situ molecular collection from living tissue; high-fidelity directed transfer and spatial decoding then recover native spatial transcriptomic information. By coupling in situ molecular capture in living tissue with subsequent molecular and phenotypic characterization, LIFE-seq aims to substantially advance our understanding of physiological and pathological processes in organs and tissues while preserving cell viability and function. In addition, spatiotemporal transcriptomic analysis in a mouse model of traumatic brain injury (TBI) revealed spatiotemporal pathological heterogeneity within the injured microenvironment. LIFE-seq thus establishes an approach for spatiotemporal transcriptomic analysis of living tissue with minimal perturbation, offering a way to resolve complex biological processes in space and advance our understanding of cellular dynamics within living tissue.

## Results

### Overview of the LIFE-seq workflow

The LIFE-seq workflow is outlined in Fig. 1. First, a nanoforest array non-destructively samples the tissue surface to obtain a complete molecular replica. In parallel, inspired by elegant designs such as Decoder-seq, we use a microfluidics-assisted barcoding strategy to fabricate spatially defined DNA capture arrays ^19^. To improve spatial barcode fidelity and assembly flexibility, we use three-segment rather than conventional two-segment probe assembly. The first segment is an initial probe containing a polymerase chain reaction (PCR) handle, which is covalently attached to an amine-functionalized glass slide. A pair of microfluidic chips (Supplementary Fig. S1a) is then placed sequentially on the slide in orthogonal orientations to deliver 50 barcode A solutions and 50 barcode B solutions. Precise fluid control, covalent coupling and two sequential DNA ligation reactions generate a DNA probe array with defined spatial coordinates at the channel intersections. As shown at the bottom of Fig. 1, the second segment contains an eight-base spatial index (Barcode A) flanked by linker sequences (Linker 1). The third segment contains a complementary linker (Linker 2), an eight-base spatial index (Barcode B), a unique molecular identifier (UMI) to remove amplification bias, and a poly(T) tract for messenger RNA (mRNA) capture. This sequential three-segment assembly yielded a significantly higher proportion of reads with valid barcodes than the two-segment strategy (P < 0.0001; Supplementary Fig. S1b, c).

**Fig 1.**
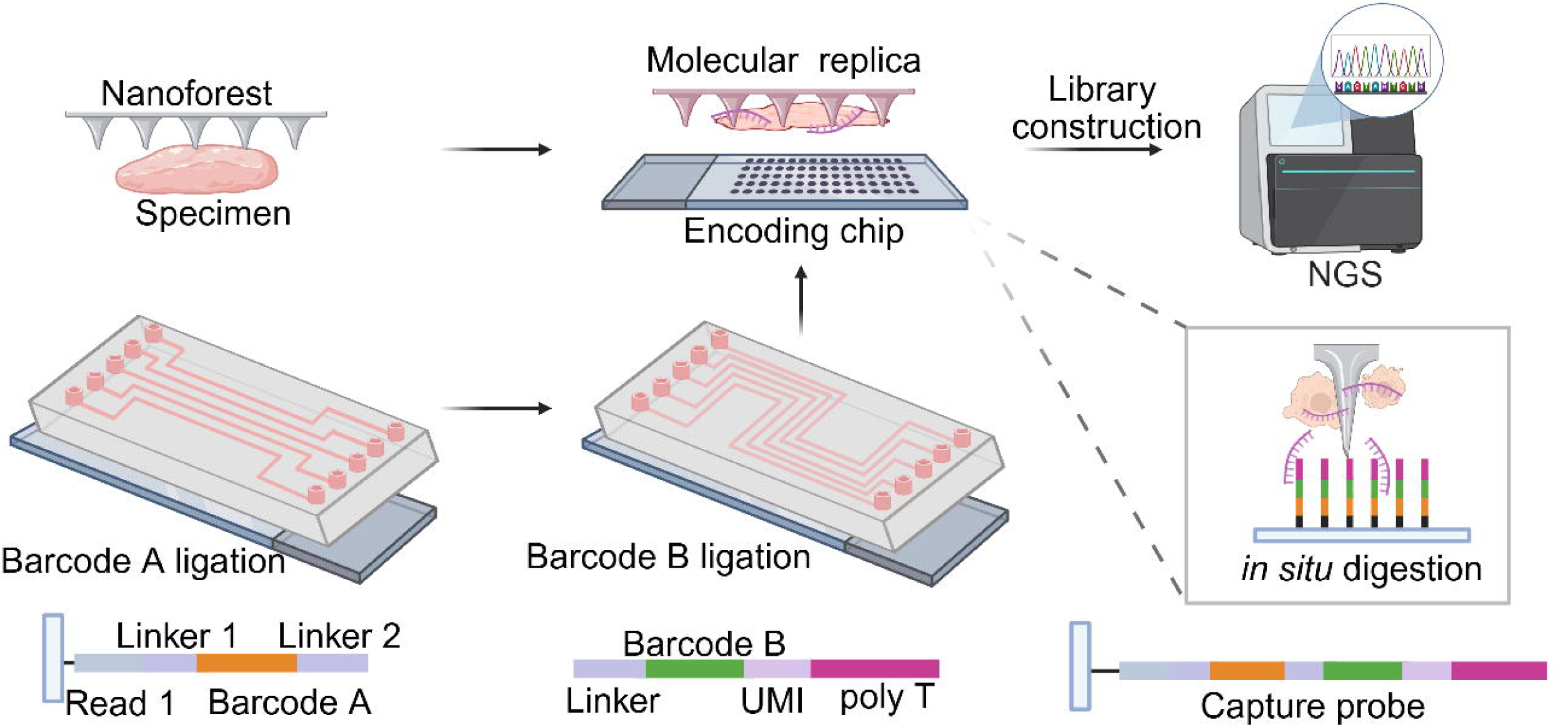
Overview of LIFE-seq. Schematic workflow of LIFE-seq, illustrating non-destructive tissue sampling via nanoforest arrays to generate molecular replicas, microfluidics-assisted orthogonal coordinate barcoding via three-segment probe assembly to fabricate high-density spatial encoding chips, alignment of the tissue-loaded nanoforest with the encoding chip for *in situ* digestion and vertical mRNA capture, and downstream next-generation sequencing (NGS) for spatial transcriptomic reconstruction.

The nanoforest bearing the tissue molecular replica is precisely aligned with the completed encoding chip, and an in situ digestion solution is introduced. Enzymatic digestion releases mRNA locally from the captured tissue, preserving its spatial origin, and the released molecules are immediately captured by poly(T) probes on the chip surface. Standard library preparation, including in situ reverse transcription, second-strand synthesis, complementary DNA (cDNA) amplification and transposase-mediated fragmentation (Supplementary Fig. S1d), is followed by next-generation sequencing (NGS). Reads containing spatial barcodes and transcript sequences are then extracted and aligned to generate high-fidelity, high-resolution spatiotemporal transcriptomic maps.

### High-sensitivity spatial transcriptomic profiling via directed molecular transfer

To establish a technical baseline for high-fidelity spatial transfer, we first developed and optimized an efficient vertical transfer system using planar glass slides (LIFE-seq-Flat; Fig. 2a). Conventional direct tissue mounting, as used in Decoder-seq and MAGIC-seq, typically requires optimization of permeabilization conditions ^19^. The mRNA released during permeabilization diffuses isotropically in solution, allowing lateral signal spread ^20^. By contrast, planar transfer avoids lengthy permeabilization optimization and vertically confines mRNA release and transport during in situ digestion. This directional transfer markedly reduces lateral diffusion and resolves tissue structures with sharper boundaries (Fig. 2b). We also tested concentration gradients of sodium dodecyl sulfate (SDS) and proteinase K (PKD) to optimize digestion, establishing 0.5% SDS as the optimal concentration (Supplementary Figs. S2 and S3). As a proof of concept, 50-μm-wide microfluidic channels were used to construct an encoding array containing 2,500 spots. Fluorescence imaging confirmed the positional accuracy and uniformity of the capture regions (Fig. 2c, d).

**Fig 2.**
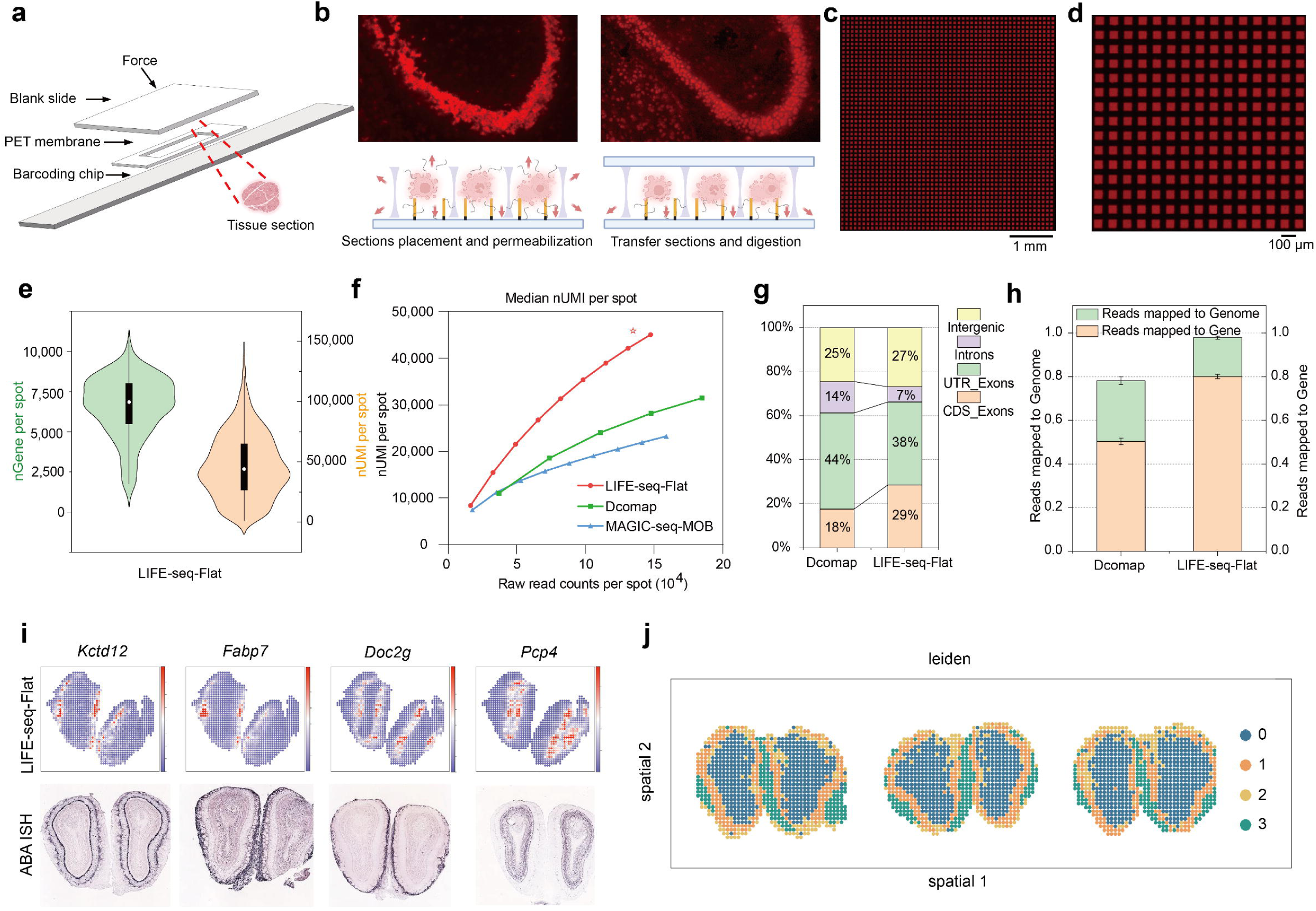
High-sensitivity spatial transcriptomic profiling via directed molecular transfer. **a,** Schematic illustrating the assembly and workflow of spatial transcriptomic profiling via planar transfer (LIFE-seq-Flat). **b,** Top: fluorescence images of captured transcripts; bottom: schematic comparison of mRNA release and capture dynamics between conventional direct tissue mounting with permeabilization (left) and directed vertical transfer with *in situ* digestion (right). **c, d,** Fluorescence microscopy images of the 50 × 50 spatially barcoded array (50-μm spot size) at low (**c**) and high (**d**) magnifications. **e,** Violin plots showing the distribution of unique genes (nGene, left) and unique molecular identifiers (nUMI, right) detected per spot in the mouse olfactory bulb (MOB) using LIFE-seq-Flat. **f,** Sequencing saturation curves plotting median UMI counts per spot against mean read counts per spot for LIFE-seq-Flat compared with direct tissue mounting and MAGIC-seq. **g,** Stacked bar chart comparing the genomic distribution of sequencing reads between direct mounting and LIFE-seq-Flat. h, Comparison of Reads mapped to genome and Reads mapped to gene between direct tissue mounting and LIFE-seq-Flat, representing the proportions of reads aligned to the reference genome and assigned to annotated genes, respectively. **i,** Spatial expression maps of four representative MOB marker genes generated by LIFE-seq-Flat (top) alongside corresponding reference *in situ* hybridization (ISH) images from the Allen Brain Atlas (bottom). **j,** Unsupervised spatial clustering (Leiden) of the MOB across three independent biological replicates using LIFE-seq-Flat.

We next systematically evaluated planar transfer using the mouse olfactory bulb (MOB). At 50-μm resolution and 47% sequencing saturation, the method detected 33,545 distinct genes, with a mean of 6,557 genes and 41,146 UMIs per spot. The distributions shown in the violin plots demonstrated high transcript capture efficiency (Fig. 2e). Sequencing saturation curves showed that LIFE-seq-Flat consistently achieved higher median UMI counts per spot across sequencing depths than direct tissue mounting or other methods at the same resolution, including MAGIC-seq ^21^, providing the highest detection sensitivity among these methods (Fig. 2f). Planar transfer also markedly improved the genomic distribution of sequencing reads (Fig. 2g), with approximately 80% mapping to annotated gene regions (Reads mapped to gene; Fig. 2h). This increased effective gene capture was primarily attributable to the cleaner environment for in situ digestion and reverse transcription in the transfer system. Spatial expression patterns of four representative MOB marker genes closely matched the in situ hybridization (ISH) reference data from the Allen Brain Atlas (ABA; Fig. 2i). Unsupervised spatial clustering identified four clusters that aligned with the distinct anatomical layers of the olfactory bulb. The highly concordant clustering patterns across three independent biological replicates demonstrated the reproducibility and spatial accuracy of vertical molecular transfer (Fig. 2j). These core performance results established a methodological foundation for integrating nanoforest arrays to profile more complex tissues.

### Non-destructive nanoforest sampling enables high-fidelity spatial transcriptomic mapping

Having established a sensitive planar vertical transfer system, we evaluated nanoforest arrays for non-destructive molecular sampling in spatial transcriptomics. Dense, disordered silica nanoneedle arrays were etched into 5 × 5 mm pure quartz substrates. The nanoneedles had a mean height of approximately 2 μm, a tip diameter of approximately 50 nm and a base diameter of 200–300 nm, forming a densely packed, forest-like three-dimensional structure with a high specific surface area (Supplementary Fig. S4). The workflow and practical procedure for non-destructive sampling of frozen mouse brain tissue are shown in Fig. 3a, b. Scanning electron microscopy (SEM) revealed the surface microstructure of the bare nanoforest and a distinct boundary of biomolecular adsorption after tissue sampling (Fig. 3c). To assess capture efficiency, we performed propidium iodide (PI) fluorescence imaging and capture-area quantification on tissue sections, planar-glass controls (Flat) and nanoforest replicas (Replica). A bright-field image of an intact mouse hemibrain served as a morphological reference. The molecular replicas faithfully reproduced the anatomical outlines of the original brain tissue, with a captured-area fraction comparable to that of intact sections, whereas planar-glass controls captured almost no detectable tissue material (P < 0.0001; Fig. 3d, e). Across successive sampling runs, SEM morphology and PI-positive replica area remained highly consistent (Supplementary Fig. S5), demonstrating the structural stability of the arrays and their capacity for repeated sampling.

**Fig 3.**
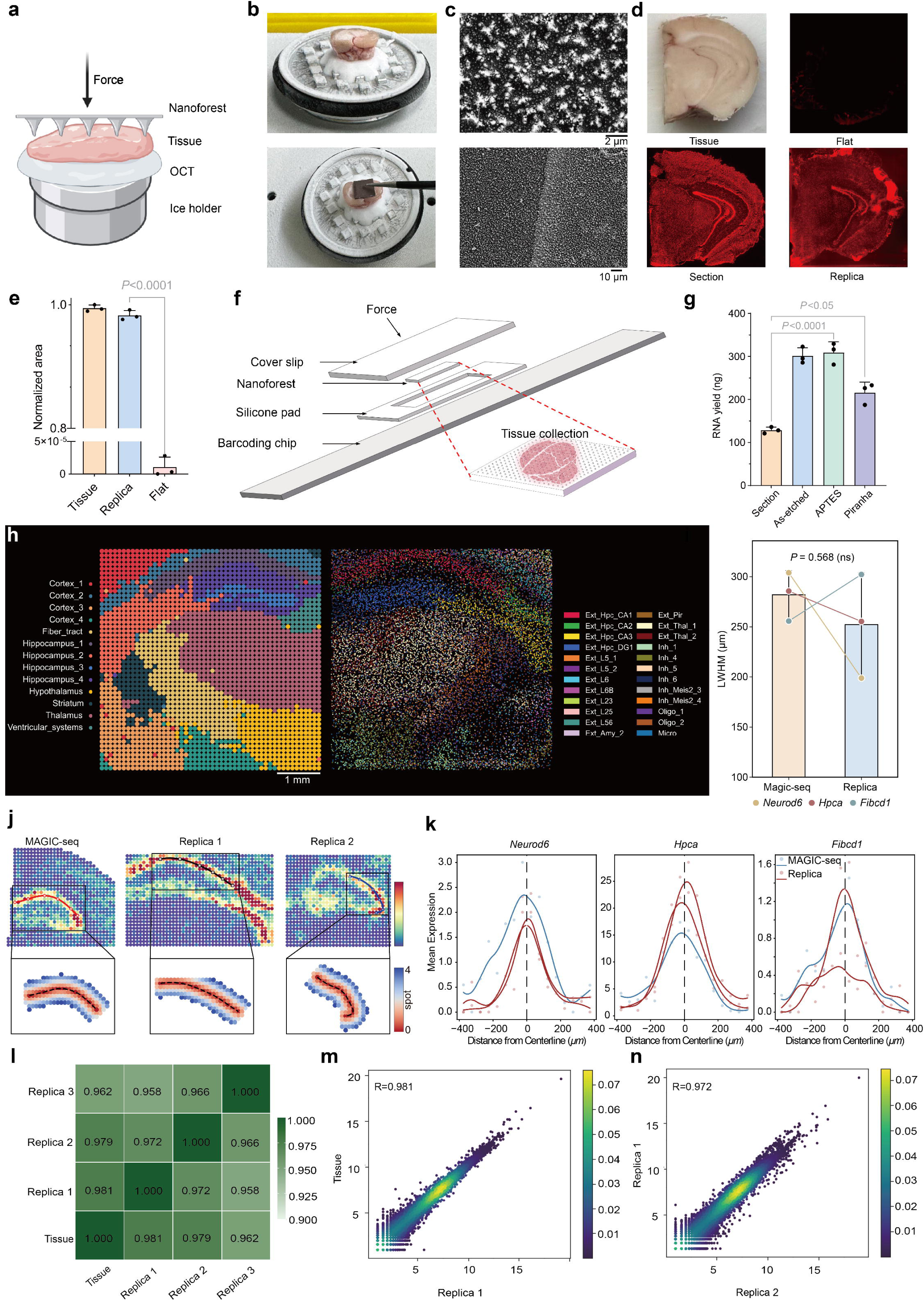
Non-destructive nanoforest sampling enables high-fidelity spatial transcriptomic mapping. **a,** Schematic of the non-destructive sampling process on frozen tissue using the nanoforest array. **b,** Photographs illustrating the practical operation of nanoforest sampling on frozen mouse brain tissue. **c,** Scanning electron microscopy (SEM) images comparing the bare nanoforest surface (top) and the boundary region after tissue sampling (bottom). **d,** Bright-field photograph of an intact mouse hemibrain (top left) and propidium iodide (PI) fluorescence staining images of the planar slide control (Flat, top right), native tissue section (Section, bottom left), and nanoforest replica (Replica, bottom right), where the Section and Replica groups show nearly identical patterns and the Flat group exhibits negligible fluorescence. **e,** Bar plot quantifying the captured PI fluorescence area across the groups shown in **d**. Data are presented as mean ± s.e.m. (*n* = 3 repeated preparations from the same tissue per condition, *P* < 0.0001, one-way ANOVA with Tukey’s multiple-comparisons test). **f,** Schematic workflow of directional vertical transfer from the tissue-loaded nanoforest to the barcoding chip, utilizing a hollow silicone pad matching the thickness of the nanoforest substrate. **g,** Total RNA recovered from 10-μm brain tissue sections (Section) and as-etched, piranha-treated and APTES-modified nanoforest arrays. Data are presented as mean ± s.e.m. (n = 3 repeated preparations from the same tissue per condition). *P* < 0.0001, Section versus APTES; *P* < 0.05, Section versus Piranha; one-way ANOVA with Tukey’s multiple-comparisons test. **h,** Spatial transcriptomic mapping of the mouse hemibrain from the nanoforest Replica group, showing unsupervised clustering annotated into 13 anatomical structures (left) and spatial mapping of deconvolved single-cell types (right). Scale bar, 1 mm. **i,** Comparison of left-width at half-maximum (LWHM) diffusion distances for *Neurod6*, *Hpca* and *Fibcd1* between MAGIC-seq and LIFE-seq replicas. Data are presented as mean ± 95% confidence interval (CI). P > 0.05, paired t-test. **j,** Spatial visualization of the region-restricted gene *Hpca* across MAGIC-seq, Replica 1, and Replica 2, highlighting the fitted boundary line along the peak expression contour and the normal vectors traversing 4 spots outward (bottom insets). **k,** Smoothed LWHM diffusion curves showing the mean expression profiles of *Neurod6*, *Hpca*, and *Fibcd1* across the boundary in MAGIC-seq, Replica 1, and Replica 2. **l,** Correlation heatmap matrix comparing global transcriptomic profiles among the native tissue section (Tissue) and three independent nanoforest replicas (Replica 1–3). **m, n,** Scatter plots showing the global transcriptomic correlation between Replica 1 and Tissue (**m**, R = 0.981) and between Replica 1 and Replica 2 (**n**, R = 0.972).

To transfer transcripts efficiently and faithfully from the nanoforest surface to the underlying spatial capture chip, we developed a multilayer vertical transfer device with a sandwich configuration (Fig. 3f). A hollow silicone spacer of uniform height constrained the position of the tissue-bearing nanoforest and aligned it vertically with the encoding chip below. Combined with in situ tissue digestion, this arrangement enabled efficient enzymatic release from the captured tissue and directed the released mRNA vertically toward the capture surface. We compared as-etched arrays with arrays hydroxylated using piranha solution or amine-functionalized with (3-aminopropyl) triethoxysilane (APTES) to assess the effects of surface treatment on tissue RNA capture (Fig. 3g). Surface functionalization increased the polarity and electrostatic adsorption capacity of the nanoforest, which recovered significantly more total RNA than tissue sections (P < 0.0001). A single sampling with an APTES-modified array recovered approximately 309 ng of RNA, compared with a mean of 128 ng extracted from a complete 10-μm tissue section (P < 0.001), providing sufficient input for downstream high-throughput spatial transcriptomic sequencing.

We then validated the complete LIFE-seq workflow by sampling frozen mouse hemibrains with nanoforest arrays. Spatial expression maps from the Replica samples resolved the complex spatial heterogeneity of brain tissue. The captured region was partitioned into 13 clusters with detailed anatomical annotations, and single-cell deconvolution further verified the spatial distribution of cell types (Fig. 3h). To quantify lateral signal spread during transfer, we selected three highly expressed hippocampal marker genes, *Neurod6*, *Hpca* and *Fibcd1*, and analyzed a published MAGIC-seq dataset ^21^ and two nanoforest replicas (Replica 1 and Replica 2). We defined contours from regions of high gene expression and extracted expression profiles spanning four spots along the outward normals. We then calculated the left-width at half-maximum (LWHM) of these profiles (Fig. 3i–k). The smoothed profiles showed similar trends for all three markers in the Replica and MAGIC-seq samples (Fig. 3j, k), and LWHM distances did not differ significantly between the two methods (P > 0.05; Fig. 3i), demonstrating that LIFE-seq preserves spatial resolution with high fidelity. Correlation analyses further confirmed the reproducibility of the transcriptomic data (Fig. 3l–n and Supplementary Fig. S6).

The inexpensive orthogonal barcoding chips and low-cost nanoneedle arrays, which can be fabricated at scale, also make LIFE-seq highly cost-effective (Supplementary Table 1). Together, these technical advances establish the methodological basis for spatiotemporal transcriptomic analysis of living tissue using LIFE-seq.

### LIFE-seq reproduces spatial transcriptomic features of traumatic brain injury

TBI is accompanied by complex, spatially heterogeneous neuroinflammation and tissue remodeling^22^. However, cellular interactions and microenvironmental changes after TBI remain relatively underexplored at the spatial level. The superficial location of cortical TBI lesions provides an ideal setting for in situ molecular profiling. We therefore applied LIFE-seq to frozen brain tissue from mice with TBI to validate its ability to capture pathological microenvironments, establishing a methodological foundation for subsequent in situ spatiotemporal monitoring in living animals.

We compared four sample groups: sham tissue sections (Sham-section), sham nanoforest replicas (Sham-replica), TBI tissue sections (TBI-section) and TBI nanoforest replicas (TBI-replica). Unsupervised spatial clustering and uniform manifold approximation and projection (UMAP) consistently resolved 12 major cellular or anatomical clusters (clusters 0–11) across the four groups, despite subtle anatomical differences among tissue samples. Cluster 3 corresponded to the TBI-specific lesion core in both TBI-section and TBI-replica samples (Fig. 4a–c). The distributions of genes and UMIs detected per spot were broadly consistent across the four groups (Fig. 4d). Nanoforest replicas showed high global transcriptomic correlations with intact tissue sections under both sham (R = 0.977) and TBI (R = 0.982) conditions (Fig. 4e, f). Biological replicates within each TBI group were also highly concordant (TBI-section, R = 0.990; TBI-replica, R = 0.983; Extended Data Fig. 1a, b). Gene detection probability curves and transcript length biases closely matched between sections and replicas (Extended Data Fig. 1c–e). For the hippocampal markers *Neurod6*, *Hpca* and *Fibcd1*, expression profiles measured along outward normals spanning six spot positions from the fitted boundary showed similar trends in Sham-section and Sham-replica samples. LWHM distances did not differ significantly between the two groups (P = 0.465; Extended Data Fig. 1f–h), further demonstrating the spatial fidelity of nanoforest replicas.

**Fig 4.**
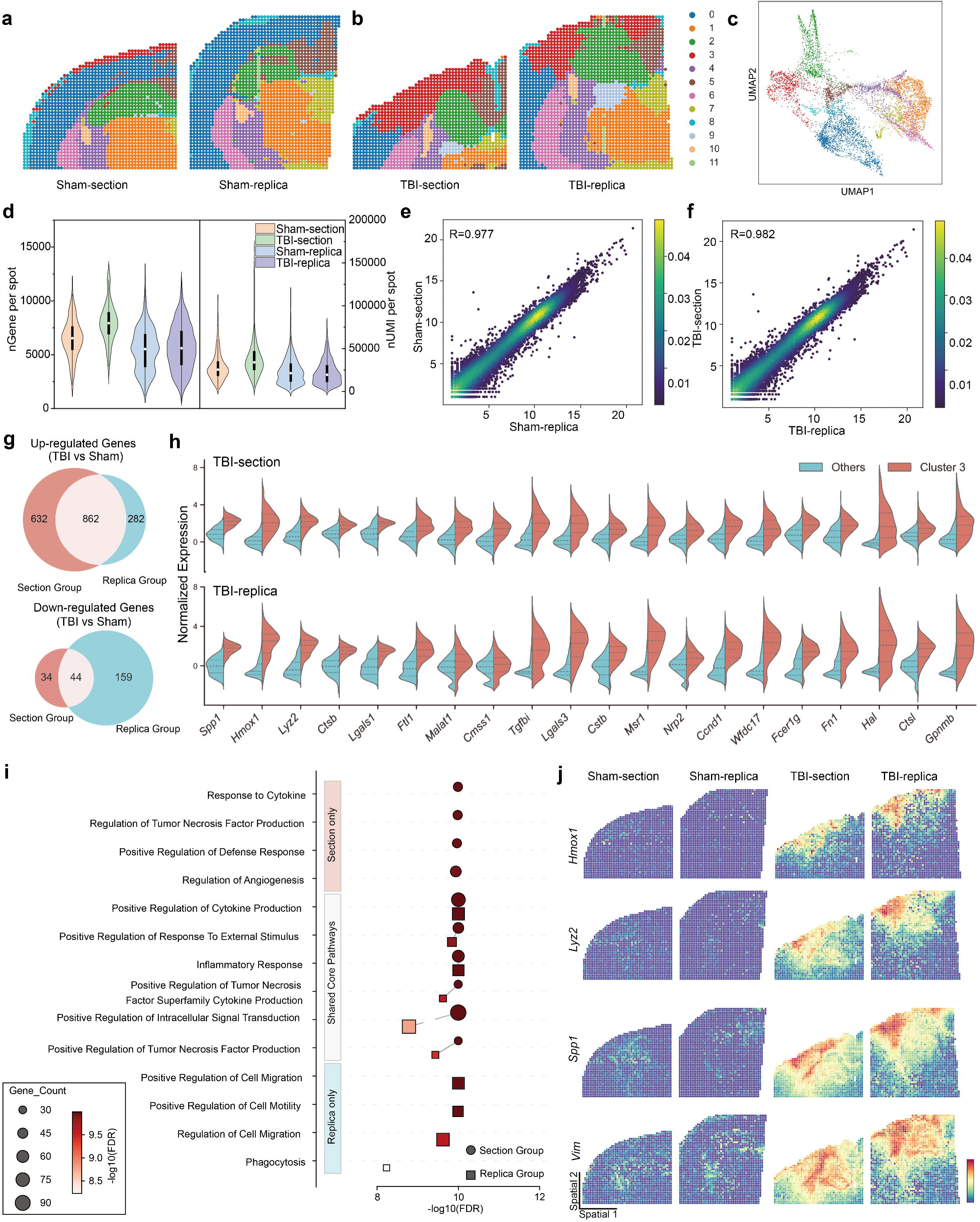
LIFE-seq reproduces spatial transcriptomic features of traumatic brain injury. **a-c**, Spatial clustering maps (**a**, **b**) and the corresponding uniform manifold approximation and projection (UMAP) embeddings (**c**) of four groups: Sham-section, Sham-replica, TBI-section, and TBI-replica. **d,** Violin plots showing the number of unique genes and unique molecular identifiers (UMIs) detected per spot across the four groups. **e, f,** Scatter plots illustrating the global transcriptomic correlations between the Section and Replica groups under Sham (**e**, *R* = 0.977) and TBI (**f**, *R* = 0.982) conditions. **g,** Venn diagrams showing the overlap of significantly upregulated (top) and downregulated (bottom) differentially expressed genes (DEGs) calculated from the TBI versus Sham comparisons in the Section and Replica groups. **h,** Split violin plots comparing the expression of the top 20 upregulated genes identified in cluster 3 of TBI-section between cluster 3 and other clusters in the TBI-section and TBI-replica groups. **i,** Comparison of the top 10 enriched Gene Ontology (GO) Biological Process (BP) terms for the upregulated DEGs in the Section and Replica groups. Six overlapping terms are highlighted. **j,** Spatial expression maps of four representative marker genes (*Vim*, *Spp1*, *Lyz2*, *Hmox1*) across the four groups. Elevated expression is highly localized to the lesion core (cluster 3) and the surrounding penumbra exclusively in the TBI groups.

We next assessed the ability of LIFE-seq to capture differentially expressed genes (DEGs) associated with pathology. MA plots and Venn diagrams showed substantial overlap between injury-responsive DEGs identified in sections and replicas (TBI versus sham; Extended Data Fig. 2a, b), including 862 commonly upregulated and 44 commonly downregulated genes (Fig. 4g). Log fold changes (logFC) were positively correlated between the two preparations (R = 0.6), confirming their concordance in capturing expression changes (Extended Data Fig. 2c). Within the lesion core, the top 20 significantly upregulated genes identified in cluster 3 of TBI-section, including *Spp1*, *Hmox1*, *Lyz2*, *Ctsb* and *Lgals3*, were expressed at significantly higher levels than in other clusters. The same genes showed a consistent pattern of high, cluster-specific expression in cluster 3 of TBI-replica (Fig. 4h). To examine the biological functions of these injury-associated genes, we performed Gene Ontology (GO) enrichment analysis of the upregulated DEGs in each preparation. Among the top 10 biological process (BP) terms, sections and replicas shared six terms closely related to acute brain injury, including positive regulation of cytokine production, inflammatory response, regulation of tumor necrosis factor (TNF) superfamily cytokine production and positive regulation of intracellular signal transduction. Replicas additionally showed enrichment of injury-response processes such as cell migration and phagocytosis (Fig. 4i). These overlapping terms captured the characteristic inflammatory and immune cascades following TBI. This concordance extended to cellular component (CC) and molecular function (MF) terms: among the top 10 terms in each category, the two preparations shared nine CC terms, including collagen-containing extracellular matrix and phagocytic vesicle, and eight MF terms, including cytokine receptor activity and Toll-like receptor binding (Extended Data Fig. 2d, e). Together, these results show that nanoforest replicas reproduce the functional enrichment patterns associated with TBI.

Finally, we examined four established pathological marker genes involved in neuroinflammation, oxidative stress and macrophage/microglial infiltration after TBI: *Vim*, a marker of astrocyte activation; *Spp1*, a microglial inflammatory-response gene; *Lyz2*, a marker of myeloid cell infiltration; and *Hmox1*, a regulator of the oxidative stress response. Spatial mapping showed generally low expression of these genes in sham sections and replicas, but strong local expression in the lesion core (cluster 3) and surrounding penumbra after TBI. Their spatial distributions and expression gradients closely matched between TBI-section and TBI-replica samples (Fig. 4j). Other anatomical markers, including *Nptxr*, *Camk2n1*, *Mbp* and *Snap25*, also retained highly consistent spatial distributions across the four groups (Supplementary Fig. S7).

Together, these results show that LIFE-seq faithfully reconstructs the fine anatomy of the brain after TBI and identifies injury-specific microenvironments and spatial expression gradients characterized by inflammatory, immune and phagocytic processes. They demonstrate the ability of nanoforest replicas to reproduce pathological features in a complex disease model and establish a foundation for subsequent in situ spatiotemporal sampling in living animals.

### LIFE-seq enables longitudinal profiling of cortical injury responses

We next used LIFE-seq to track transcriptional changes following cortical injury in the same animal. In a mouse model of focal cortical stab wound injury (SWI), we performed in situ sampling before injury (D0) and on D1 and D2. Together with terminal tissue analysis on D3, these samples formed a spatial transcriptomic series spanning four time points (Fig. 5a). We obtained 10 spatial expression datasets, with high numbers of genes and transcripts detected at each time point. Global expression correlations between samples ranged from 0.910 to 0.986 (Supplementary Fig. S8).

**Fig 5.**
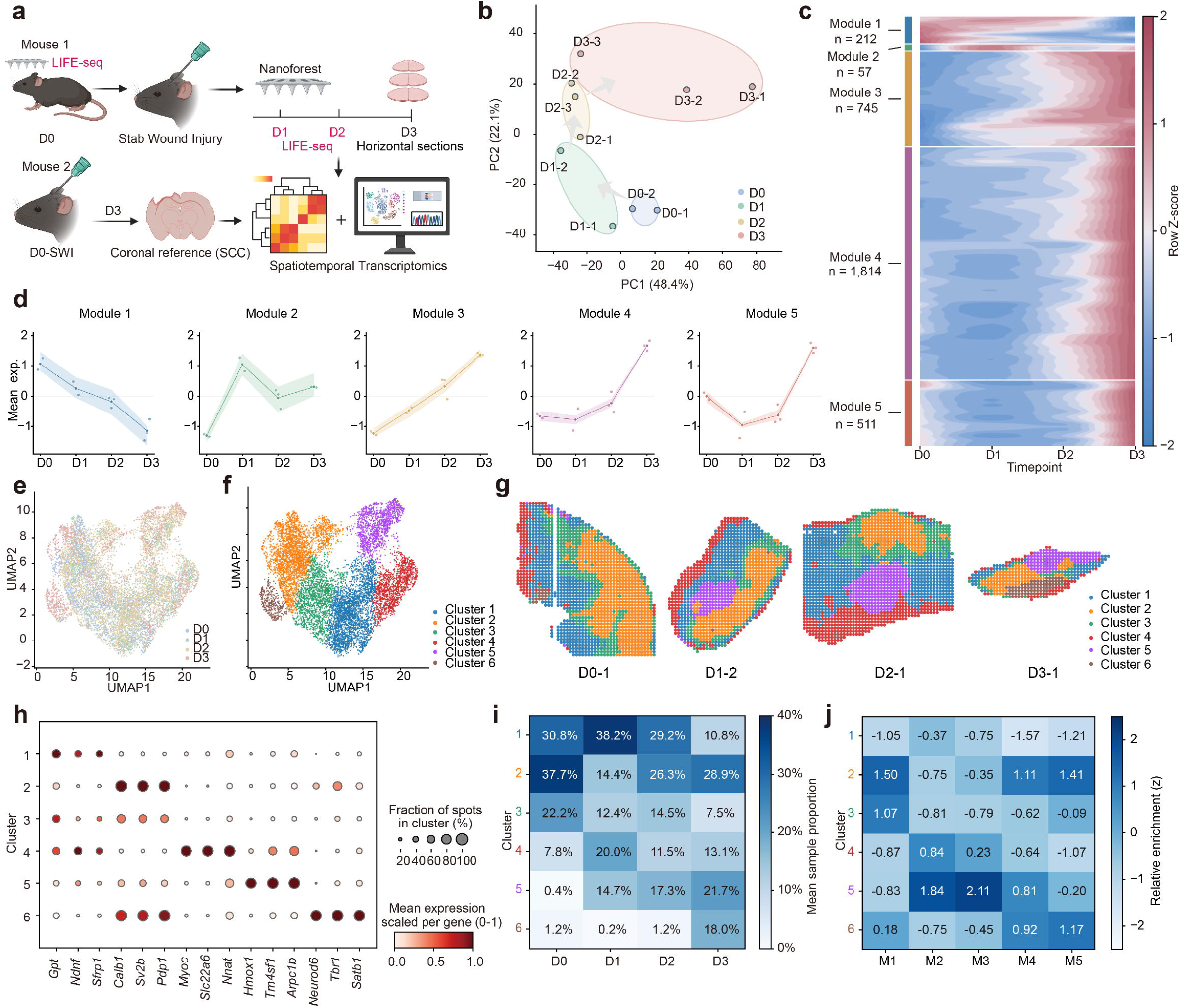
LIFE-seq enables longitudinal profiling of cortical injury responses. **a,** Experimental design. In the same mouse, LIFE-seq sampling was performed before cortical stab injury at D0 and after injury at D1 and D2. At D3, endpoint spatial transcriptomes were obtained by direct mounting of horizontal cryosections from this mouse after euthanasia. D0, D1, D2 and D3 comprised two, two, three and three samples, respectively, yielding ten samples from the same animal. A second mouse subjected to SWI at D0 was euthanized at D3 to provide the SWI coronal control (SCC). **b,** Sample-level principal component analysis. Each point represents one sample, colored by time point. Axis labels indicate the percentage of variance explained. Shaded ellipses guide time-point grouping, and arrows indicate temporal order. **c,** Heatmap of 3,339 temporally varying genes grouped into five patterns according to their standardized expression trajectories across four time points. Colors indicate gene-wise Z-scores, and n denotes the number of genes in each pattern. Interpolation and smoothing were applied for heatmap display. **d,** Mean standardized expression trajectories of the five temporal patterns. Lines represent the mean across all member genes, small points show sample-level pattern means, and shading indicates the 25th–75th percentiles of member-gene Z-scores. **e, f,** Joint UMAP of all 10,270 spatial spots, colored by time point (e) or expression cluster (f). **g,** Spatial distributions of the six expression clusters in D0-1, D1-2, D2-1 and D3-1. **h,** Representative gene expression across clusters. Dot size indicates the percentage of spots detecting each gene within a cluster; color indicates mean expression scaled to 0–1 for each gene. **i,** Mean spot proportions of the six clusters at each time point. Proportions were calculated within each sample and averaged with equal sample weights. **j,** Relative scores of the five temporal patterns across the six expression clusters. Mean expression of pattern members was adjusted for the overall background within each sample, averaged with equal sample weights and Z-standardized across clusters separately for each pattern.

Analysis across consecutive time points revealed expression responses with distinct temporal courses after cortical injury. Principal component analysis (PCA) showed stage-dependent shifts in the overall transcriptional state of the samples, reflecting continuing changes after injury (Fig. 5b). We identified five temporal patterns from the expression trajectories of 3,339 time-associated genes (Fig. 5c, d). Pattern 2 genes, including *Lcn2*, *Gfap* and *Serpina3n*, peaked at D1 and subsequently declined, indicating a transient early response. In pattern 3, *Lyz2*, *Spp1* and *C1qb* increased progressively from D0 to D3, showing a sustained rise in expression. Whereas both patterns began to increase soon after injury, the increase in pattern 4 occurred mainly at D3, indicating a later response. Pattern 1 declined continuously, whereas pattern 5 decreased early and rebounded at D3, revealing trajectories that a single endpoint comparison would not readily distinguish. These trajectories separated post-injury transcriptional changes into transient early responses, sustained increases and late changes, revealing distinct stages of the injury response.

Transcriptional changes associated with glial responses emerged early after injury. Genes upregulated at D1 relative to D0 included *Lcn2*, *Gfap* and *Serpina3n*, a characteristic expression signature of the glial response to cortical SWI ^23^. *Hmox1* and *Ifitm3* were also upregulated, reflecting early stress and inflammatory responses (Supplementary Fig. S9a). More extensive expression changes were observed at D3 relative to D0 (Supplementary Fig. S9d). Functional enrichment linked genes upregulated at D1 mainly to inflammatory responses, cell migration, chemotaxis and regulation of angiogenesis. At D3, enriched functions additionally included leukocyte activation, innate immune responses and regulation of cytokine production (Supplementary Fig. S9e,f). This temporal course revealed the continuing development of the local injury response. Inflammation initiated early after injury persisted to D3 and was accompanied by broader immune-related expression changes.

Joint clustering of all spatial spots identified six expression clusters with localized distributions in the tissue (Fig. 5e–g and Supplementary Fig. S10a,b). Cluster 5, characterized by higher expression of *Hmox1*, *Tm4sf1* and *Arpc1b*, formed local aggregates in post-injury samples. It accounted for a higher proportion of sampled spots than at D0 (Fig. 5g–i and Supplementary Fig. S10c). This cluster had the highest relative expression scores for both the transient early pattern 2 and the progressively increasing pattern 3. Pattern 5, which declined early and subsequently rebounded, scored higher in Clusters 2 and 6 (Fig. 5j). These results linked post-injury expression trajectories to local expression features in tissue, demonstrating the ability of LIFE-seq to resolve spatiotemporal injury responses within the same animal.

### LIFE-seq enables characterization of cellular heterogeneity and spatial remodeling following cortical injury

To resolve cell composition in the injured region, we applied robust cell type decomposition (RCTD) using a published single-cell reference from SWI cortex ^23^. We analyzed the terminal coronal section (SWI coronal control, SCC) and sample D2-1 (Fig. 6a and Extended Data Fig. 4). The C1 expression region in SCC corresponded to the histological injury site (Fig. 6b and Extended Data Fig. 4c). Mapping the SCC expression regions onto D2-1 identified a C1-like region, around which we partitioned the surrounding tissue by distance from C1 (Fig. 6c–f). Macrophage/monocyte-associated signals were concentrated in C1. Astrocyte group 13 (13_Astrocytes) had a higher RNA contribution around C1, whereas astrocyte group 6 had a higher RNA contribution farther away (Extended Data Fig. 4d).

**Fig 6.**
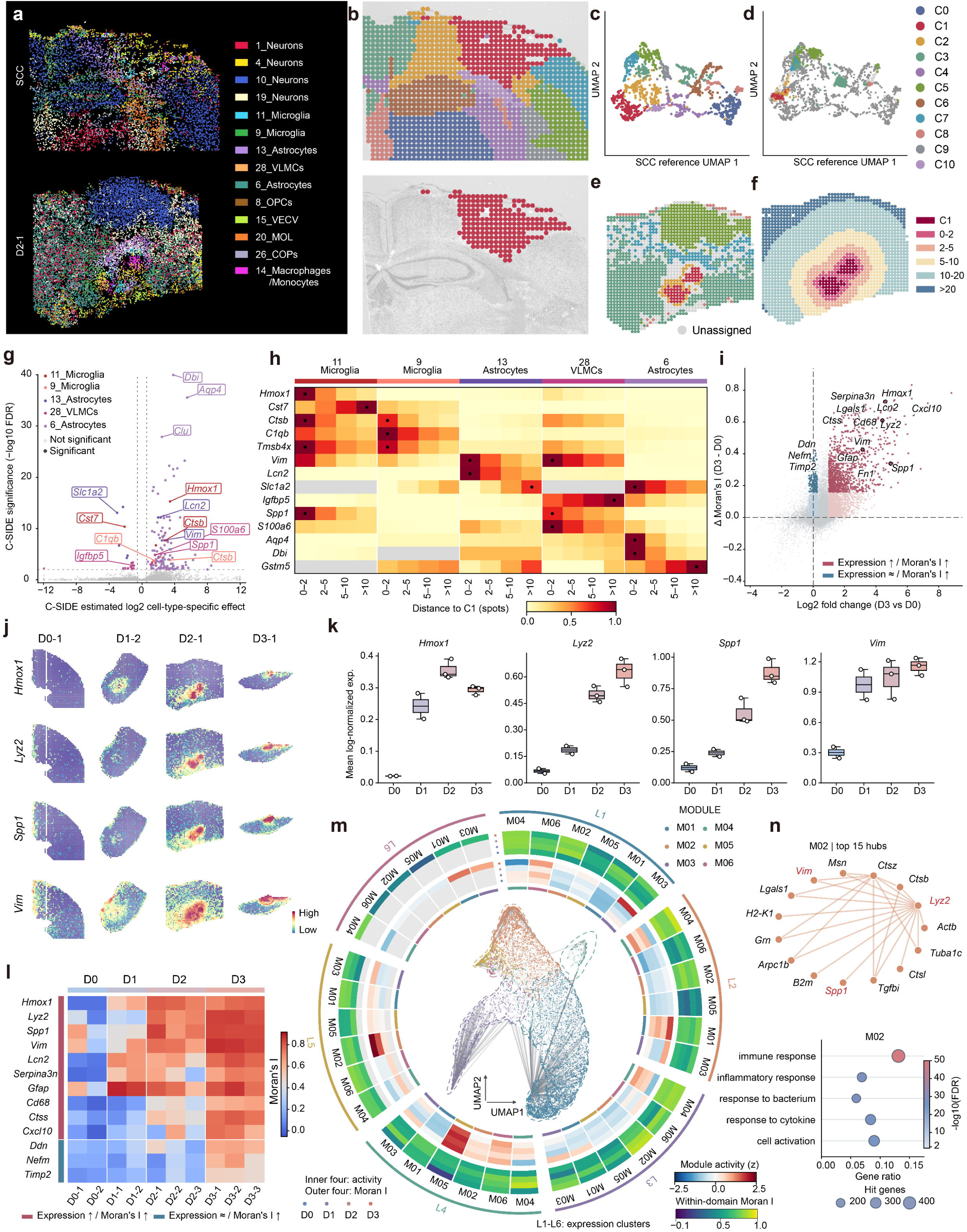
LIFE-seq enables characterization of cellular heterogeneity and spatial remodeling following cortical injury. **a,** Spatial mapping of cell types in the endpoint coronal section (SCC) and D2-1 using robust cell type decomposition (RCTD) with a published single-cell reference. **b,** Spatial distribution of 11 expression clusters (C0–C10) in SCC (top) and localization of the injury-associated cluster C1 on the corresponding histological image (bottom). **c,** UMAP of the SCC reference colored by expression cluster. **d,** Projection of D2-1 spots onto the SCC reference UMAP. **e,** Spatial mapping of SCC-derived cluster annotations in D2-1, with unassigned spots shown in grey. **f,** Distance-based stratification of D2-1 spots relative to C1 (C1, 0–2, 2–5, 5–10, 10–20 and >20 spots). **g,** C-SIDE analysis showing estimated log2 cell-type-specific effects and −log10(FDR), with significant genes colored by cell type. Representative genes are labeled. **h,** Heatmap of normalized cell-type-specific expression estimates for representative genes across five cell populations and four distance intervals from C1 (0–2, 2–5, 5–10 and >10 spots). **i,** Scatter plot of log2 fold changes (D3 versus D0) against changes in spatial autocorrelation (ΔMoran’s I, D3 − D0). Red and blue indicate genes with increased expression and Moran’s I, or relatively unchanged expression with increased Moran’s I, respectively. **j,** Spatial expression maps of *Hmox1*, *Lyz2*, *Spp1* and *Vim* across four representative samples (D0-1, D1-2, D2-1 and D3-1). **k,** Box plots showing mean log-normalized expression of the four genes across D0–D3. Points represent individual samples. **l,** Heatmap of Moran’s I values for representative genes across ten samples, grouped by changes in expression and spatial autocorrelation. **m,** Circular visualization of six gene coexpression modules (M01–M06) across six spatial expression clusters (L1–L6). The central UMAP shows the gene coexpression network, while the inner and outer rings represent module activity (Z-score) and within-domain Moran’s I across D0–D3, respectively. **n,** Coexpression network of the top 15 hub genes in module M02 (top) and representative enriched Gene Ontology (GO) Biological Process (BP) terms (bottom). Bubble size indicates gene count, and color indicates −log10(FDR). D0–D2 samples were collected in vivo using LIFE-seq, whereas D3 samples were obtained by conventional tissue-section mounting after euthanasia. All ten samples originated from the same mouse.

Cell type-specific inference of differential expression (C-SIDE) further revealed spatial variation in gene expression within individual cell types. After accounting for cell composition, *Hmox1* and *Spp1* expression in 11_Microglia increased near the injury (Fig. 6g, h). Although the RNA contribution of 13_Astrocytes was higher around C1, their reactivity-associated genes showed stronger expression closer to C1. This distinction revealed local changes in cell state beyond differences in cell distribution. Astrocytes near the injury had a stronger reactive expression profile and lower expression of the glutamate transporter gene *Slc1a2*. Together with gene expression distributions in the single-cell reference, these findings indicated that perilesional tissue differences involved both cell-population distributions and within-cell-type expression states (Extended Data Fig. 4e).

Changes in gene expression after injury were accompanied by changes in spatial distribution. From D0 to D3, *Hmox1*, *Lyz2*, *Spp1* and *Vim* expression increased, forming more distinct regions of high expression. These changes were accompanied by increases in Moran’s I (Fig. 6i–l and Extended Data Fig. 3a, c). Other genes showed little change in overall expression but similarly increased spatial autocorrelation. For example, *Ddn* and *Timp2* developed more distinct regional expression differences after injury, which were less apparent in tissue-averaged expression (Extended Data Fig. 3b). Functional enrichment linked genes with increases in both expression and spatial aggregation primarily to inflammatory and immune responses. Genes with increased spatial autocorrelation but little overall expression change were associated with synapse organization, regulation of synaptic transmission and neuron projection extension (Extended Data Fig. 3d).

Coexpression analysis across time points identified six gene modules whose activity and spatial autocorrelation differed among expression regions (Fig. 6m). M02 was enriched for processes including immune responses, inflammatory responses and responses to cytokines (Fig. 6n). Spatial mapping showed M02 activity increasing progressively from a low level at D0. Prominent regions of high activity appeared at D2 and D3 (Supplementary Fig. S11). In D2-1, these regions corresponded to C1, and module activity decreased with increasing distance from C1 (Extended Data Fig. 4f, g). This finding extended injury-proximal immune expression from individual genes to a set of genes that varied together. The localized distribution was also consistent with injury-proximal immune responses observed in studies of focal brain contusion ^24^.

These results show that cortical injury responses involve enhanced immune-related programs, spatial reorganization of neural gene expression and local, cell-type-specific expression changes. LIFE-seq links the temporal course of transcriptional responses to these local features, resolving microenvironmental changes that tissue-averaged expression cannot readily reveal. Thus, LIFE-seq extends spatial transcriptomics to longitudinal studies of the pathological microenvironment within the same animal. It provides a new way to observe how molecular responses arise, are distributed and evolve during tissue injury.

## Discussion

Biological activity is highly dynamic in space and time, and understanding its complexity requires examining it in situ within the native physiological microenvironment ^2^. Conventional tools, including bulk sequencing, fixed-tissue analyses and endpoint assays, capture only isolated snapshots. Linking molecular events across timescales to real-time gene expression and tracking cellular dynamics with minimal physiological perturbation have therefore become central goals in resolving the spatiotemporal molecular dynamics of living tissue ^11^. Emerging technologies are moving toward these goals but retain substantial limitations. Conventional ST is restricted to fixed or frozen specimens. Intravital optical imaging depends heavily on fluorescent reporters and lacks unbiased transcriptome-wide profiling ^25^, whereas microelectrodes ^26^, microdialysis ^27^, and *in vivo* sensors ^28^ are limited by narrow target coverage or insufficient spatial resolution. Metabolic labeling can record temporal information ^29^, but generally requires complex chemical or genetic modifications and ultimately relies on destructive tissue lysis at the endpoint. Existing tools therefore struggle to capture genome-wide transcriptional dynamics without bias and at high spatial resolution while maintaining tissue viability. To address these limitations, we developed LIFE-seq by integrating non-destructive in situ sampling with nanoforest arrays and microfluidic spatial combinatorial indexing, enabling detailed spatiotemporal transcriptomic analysis of living tissue.

In developing ST for living tissue, direct functionalization of microneedle surfaces with spatial barcodes is a potential strategy, but it entails complex assembly and high manufacturing costs, and its ultimate spatial resolution is constrained by needle spacing. Recent strategies that extract RNA from living tissue or fresh sections with silicon nanoneedle arrays and then hybridize spectrally encoded probes have enabled sequencing-free spatial mapping ^16, 17^. However, their reliance on predefined targets limits gene-detection throughput and unbiased discovery. LIFE-seq instead decouples molecular sampling in living tissue from subsequent spatially resolved capture. First, high-density nanoforest arrays enable high-fidelity, minimally invasive molecular replication. At only 2 μm in height, the arrays combine a high specific surface area with submicrometer resolution at the sampling interface. This avoids the superposition of signals from different depths caused by deep penetration with conventional long microneedles, preserves the original two-dimensional tissue anatomy and minimizes physiological perturbation during sampling. More importantly, minimally invasive sampling enables longitudinal analysis at multiple time points within the same individual, reducing the influence of interindividual variation on studies of tissue transcriptional dynamics. Second, the spatial mapping accuracy of LIFE-seq is no longer directly limited by the physical spacing of the sampling structures and can be further improved by optimizing the barcode density of the downstream capture chip. This standardized transfer scheme makes the system modular. It has been implemented with our microfluidically barcoded capture chips and offers the potential for compatibility with mainstream commercial 3’-end ST platforms, such as 10x Visium and Stereo-seq ^30^. In addition, mechanically constrained vertical transfer keeps mRNA release highly directional, substantially suppressing lateral diffusion and preserving molecular localization while eliminating the extensive optimization of tissue permeabilization, thereby simplifying the workflow. Finally, established, low-cost fabrication methods for nanoforest arrays substantially reduce the cost per experiment, providing a broadly applicable solution for large-scale, high-throughput spatiotemporal omics in living tissue.

Several developments could further improve LIFE-seq. First, nanoforest arrays built on rigid, planar substrates may make uneven contact with nonplanar tissue surfaces during molecular replication, reducing spatial accuracy. Fabrication on flexible substrates could improve conformal contact with complex tissue surfaces. Second, LIFE-seq is currently suited primarily to superficial lesions, such as skin lesions and superficial tumors, or physically exposed tissue planes. Directly resolving three-dimensional spatial heterogeneity within solid tumors or deeper regions of complex organs without disrupting the overall tissue architecture remains difficult. Future integration with minimally invasive clinical devices, including endoscopes ^16^, medical catheters and wearable patches, would extend LIFE-seq beyond the body surface to hollow organs, such as the intestine and respiratory tract, enabling longitudinal, non-destructive sampling of lesions in situ for early molecular screening and dynamic companion diagnostics. In addition, physical positioning alone cannot ensure exact alignment of sampling coordinates across time points. Resulting spatial offsets may cause pre-existing local heterogeneity to be misinterpreted as transcriptional dynamics. Dedicated spatial registration algorithms will therefore be needed to eliminate this bias. Finally, LIFE-seq currently captures only mRNA with poly(A) tails. Its open architecture could support integration with proteomic, metabolomic and other multi-omic analyses to more fully resolve regulatory networks in cells in their native states.

In summary, LIFE-seq is a broadly compatible, low-cost and minimally invasive spatiotemporal omics tool that enables longitudinal ST analysis of living tissue within the same individual. It offers a new approach to investigating the spatiotemporal evolution of complex biological systems in their native physiological state and provides a strong technical foundation for in situ molecular diagnosis and longitudinal disease monitoring.

## Methods

### Animals and experimental design

Male C57BL/6J mice aged 6–7 weeks were maintained with free access to food and water. All animal experiments were approved by the Animal Ethical and Welfare Committee of Zhongda Hospital, Southeast University (Approval No. 20230206002) and performed in accordance with the approved institutional guidelines. Mouse olfactory bulb and brain tissues were used for planar-transfer and nanoforest-sampling experiments. Controlled cortical impact (CCI) was used for the frozen-tissue injury comparison, whereas SWI was used for longitudinal sampling.

The CCI–sham comparison included three mice: one sham mouse provided Sham-section and Sham-replica, and two CCI mice each provided a matched section–replica pair (TBI-section 1/TBI-replica 1 and TBI-section 2/TBI-replica 2, respectively).

For CCI, mice were anesthetized with isoflurane (3–4% induction and 2% maintenance) and secured in a stereotaxic frame. A craniotomy was made over the left cortex, centered 2.5 mm posterior and 2.5 mm lateral to bregma. A 3-mm impactor delivered a single impact at 5.5 m/s, with a depth of 1.0 mm and a dwell time of 300 ms. Sham animals underwent the same anesthesia and craniotomy without impact. The wound was sutured, and mice recovered on a heated pad. Brains were collected three days after injury following deep anesthesia and euthanasia by cervical dislocation.

For SWI, the left sensorimotor cortex was exposed under isoflurane anesthesia at approximately 2.0 mm posterior and 2.0 mm lateral to bregma. A sterile 20-gauge needle was inserted vertically to a depth of approximately 1.0 mm, held for 1 min and withdrawn slowly. Two mice were used for the longitudinal and reference experiments: one underwent baseline sampling before injury and was followed through D3, and the other underwent SWI at D0 and provided a terminal coronal-section reference at D3.

### Nanoforest array fabrication and surface functionalization

Nanoforest arrays were fabricated on 5 × 5 mm pure quartz (silica, SiO2) substrates by plasma etching through a self-organized polymer nanofiber mask. After transfer of the nanostructure pattern into the quartz and removal of the residual mask, the resulting disordered nanostructures were approximately 2 μm high, with tip diameters of approximately 50 nm and base diameters of 200– 300 nm. Array morphology was examined by SEM before and after tissue sampling.

Three surface conditions were evaluated: as-etched arrays, arrays hydroxylated with piranha solution and arrays functionalized with APTES. For amination, hydroxylated arrays were incubated in 2% APTES in ethanol for 2 h at room temperature, rinsed with ethanol and water, and baked at 110 °C for 1 h. RNA recovery was compared among the surface treatments, and APTES-functionalized arrays were used for subsequent nanoforest sampling.

### Microfluidic device design and fabrication

Microfluidic master molds were fabricated by SU-8 photolithography. Polydimethylsiloxane (PDMS) base and curing agent were mixed at 10:1, degassed, cast onto the molds and cured at 80 °C for 2 h. The slabs were peeled off, trimmed and punched to form inlet and outlet ports. Each device contained 50 parallel channels, 50 μm wide and separated by 50-μm gaps.

Two PDMS devices were applied sequentially in perpendicular orientations for spatial barcoding. Each device was aligned with the capture slide and secured in a custom clamp to maintain contact and prevent leakage. The orthogonal channel intersections defined 50 × 50 capture sites, each 50 × 50 μm in area, with a 100-μm center-to-center pitch.

### Preparation of spatially barcoded capture chips

Capture slides were functionalized with a γ-glycidoxypropyltrimethoxysilane (GOPTS)– poly(amidoamine) (PAMAM) layer. Hydroxylated glass was treated with GOPTS to introduce epoxy groups, followed by coupling of generation-4 PAMAM dendrimers through a ring-opening reaction ^19^. Amino-modified Read 1 probes containing the PCR handle were immobilized at 40 μM on the resulting amine-rich surface using disuccinimidyl suberate (DSS) at room temperature for 1 h. Unreacted surface amines were blocked with succinic anhydride before washing.

Fifty Barcode A oligonucleotides and fifty Barcode B oligonucleotides were separately annealed to their complementary ligation linkers. Each channel of the first PDMS device received a mixture containing 10 μM of a distinct Barcode A and 40 U/μL T4 DNA ligase. Ligation was performed at room temperature for 2 h in a humid chamber. The device was removed, and the slide was washed with 2× saline–sodium citrate (SSC) containing 0.1% SDS, followed by 0.2× SSC and water. The second device was then applied perpendicular to the first to ligate the 50 Barcode B probes at the same concentrations and under the same conditions.

After the second ligation, complementary linkers were removed with 80 mM KOH for 30 s. The chips were rinsed with water, dried and stored protected from light until use. Each assembled probe contained two 8-base spatial indices, a 12-base UMI and a poly(T) capture tract. All oligonucleotide sequences and modifications used for chip preparation, reverse transcription, second-strand synthesis, amplification, indexing and fluorescence validation are provided in Supplementary Table 2.

### Tissue preparation and nanoforest sampling

Brains used for frozen-tissue nanoforest sampling were snap-frozen in liquid nitrogen without embedding and stored at −80 °C. Samples were equilibrated on the −20 °C cryostat stage and attached to the specimen holder using optimal cutting temperature (OCT) compound only at the base. When the exposed tissue surface began to thaw slightly on ice, a nanoforest array was placed face-down on the surface for 2–3 s and lifted off. The replica was air-dried for 1 min and rapidly frozen until processing. Adjacent 10-μm cryosections and flat glass substrates served as tissue-section and sampling controls, respectively.

The longitudinal mouse provided two, two and three in vivo nanoforest captures at D0 before SWI and at D1 and D2 after injury, respectively. At D3, the mouse was euthanized, and three endpoint samples were prepared by direct mounting of horizontal cryosections through the injury region. A second mouse underwent SWI at D0 and was euthanized at D3 to provide the SCC for histological and transcriptomic annotation. SCC was analyzed separately from the ten longitudinal captures.

In vivo sampling was performed through the cortical window under isoflurane anesthesia. The nanoforest was placed over the target cortical area before injury or over the injury region after SWI, allowed to contact the tissue for 2–3 s and retrieved to obtain a local tissue imprint. After sampling, hemostasis and wound closure were performed, and the mouse recovered on a 37 °C heating pad before being returned to its cage.

### Directed molecular transfer and mRNA capture

For LIFE-seq-Flat, 10-μm frozen sections were mounted on plain glass slides, fixed in prechilled methanol, stained with hematoxylin and eosin (H&E), and imaged. The sections were destained with 0.1 M HCl for 15 min and rinsed before transfer. A 10-μm polyethylene terephthalate (PET) spacer surrounded the capture region, and the tissue-bearing slide was aligned face-down over the barcoded surface. For nanoforest replicas, the tissue-loaded array was positioned opposite the capture chip in a sandwich assembly, using a hollow silicone spacer matched to the substrate thickness.

A 30-μL transfer mixture was prepared with 1.5 μL of 10% SDS, 1.2 μL of proteinase K (20 μg/μL), 6 μL of 5× reverse-transcription buffer and RNase-free water to volume, giving final concentrations of 0.5% SDS, 0.8 mg/mL proteinase K and 1× buffer. The mixture was applied between the donor substrate and capture chip, and the assembly was incubated at 37 °C for 30 min to release RNA and capture it on the opposing poly(T) probes. The donor substrate was then removed, and the capture chip was washed with 0.1× SSC containing RNase inhibitor before immediate reverse transcription.

Direct-mounted brain-section controls were fixed in methanol and permeabilized with 0.1% pepsin in 0.1 M HCl at 37 °C for 15 min ^21^. The capture surface was washed with 0.1× SSC before reverse transcription.

### Reverse transcription and cDNA synthesis

Captured mRNA was reverse-transcribed on the chip in a reaction containing 5% polyethylene glycol 8000 (PEG8000), 1× reverse-transcription buffer, 10 U/μL Maxima H Minus reverse transcriptase, 2 U/μL RNase inhibitor, 0.5 mM deoxynucleoside triphosphates (dNTPs), 2.5 μM template-switching oligonucleotide (TSO) and 1 mM guanosine triphosphate (GTP). The reaction was incubated at 42 °C for 2 h. After washing with 0.1× SSC, the RNA template was removed with 80 mM KOH at room temperature for 5 min, followed by additional SSC washing.

Second-strand synthesis used a 37.5-μL reaction containing 4 mM MgSO4, 1× isothermal amplification buffer, 0.32 U/μL Bst 2.0 DNA polymerase, 1 mM dNTPs and 2.5 μM second-strand primer, supplemented with RNase-free water.

Second-strand synthesis was performed at 65 °C for 15 min. After SSC and elution-buffer washing, the newly synthesized strand was released with 40 μL of 80 mM KOH for 10 min and neutralized with 6 μL of 1 M Tris buffer (pH 7.0).

For cDNA amplification, 50 μL of KAPA HotStart ReadyMix and 2 μL each of 20 μM Amp-F and Amp-R primers were added to the neutralized cDNA. PCR consisted of initial denaturation at 98 °C for 3 min; 14 cycles of 98 °C for 20 s, 65 °C for 30 s and 72 °C for 2 min; and a final extension at 72 °C for 5 min. Amplified cDNA was purified using a 0.7× magnetic-bead-to-sample ratio and quantified before library preparation.

### Library construction and sequencing

Purified cDNA was quantified using a Qubit fluorometer and assessed for fragment-size distribution. Five nanograms of cDNA was used for Tn5-mediated tagmentation with the TruePrep DNA Library Prep Kit V2 (Vazyme, TD512) at 55 °C for 10 min, followed by sample-index PCR. Libraries were purified with magnetic beads, quantified and assessed for fragment-size distribution before paired-end 150-bp Illumina sequencing. The spatial barcode pair and UMI were retained for assigning transcripts to capture positions and removing amplification duplicates.

### Microscopy and assessment of sampling performance

Nanoforest replicas, conventional sections and flat-substrate controls were stained with PI (1:100 in phosphate-buffered saline (PBS)) for 6 min at room temperature and washed with PBS. Fluorescent capture areas were quantified using ImageJ. SEM and PI imaging were also used to assess array morphology and tissue capture after repeated sampling and cleaning cycles.

For RNA-yield comparisons, total RNA was extracted from 10-μm brain sections and tissue-bearing arrays with different surface treatments using a standard TRIzol/chloroform-substitute extraction procedure. Recovered RNA was quantified using the Equalbit RNA HS Assay Kit (Vazyme, EQ211-01).

Capture-chip uniformity was assessed by hybridization with Cy3-labeled poly(A) oligonucleotides followed by fluorescence scanning. Fluorescent cDNA footprints were generated by incorporating Cy3-dCTP during reverse transcription. H&E-stained brain sections were imaged by bright-field microscopy, and the SCC image was used to relate the injury region to transcriptomic domains.

### Sequencing data processing and quality control

Raw paired-end FASTQ files were processed using custom Python scripts to extract spatial barcodes and UMIs and remove adapter- and template-switching-related sequences. All mouse datasets were aligned to the same mm39 reference genome with GENCODE vM29 annotation ^31^ using STARsolo^32^. A whitelist of the 50 × 50 barcode combinations was used for barcode assignment and UMI deduplication to generate gene-by-spot count matrices. Barcode identities were mapped to the physical capture coordinates.

Count matrices were imported into Scanpy and AnnData ^33^. For tissue sections, available histological images were registered to the capture grid to identify tissue-covered positions. Valid sampling regions in in vivo nanoforest captures were identified using the spatial distribution of UMI counts and other quality metrics, together with manual registration. Sample-level quality control considered UMI counts, detected genes and mitochondrial transcript fractions; low-quality spots and sparsely detected genes were removed. Counts were normalized to 10,000 per spot and log1p-transformed, while raw counts and native spatial coordinates were retained for count-based and spatial analyses.

### Spatial clustering and anatomical annotation

Spatial expression domains were identified by unsupervised clustering of quality-controlled spots. Highly variable genes were selected from log-normalized expression. Expression values were scaled before PCA. An expression-neighborhood graph was constructed from the principal components and used for Leiden clustering ^34^ and UMAP. Cluster markers were identified using the Wilcoxon rank-sum test.

For the longitudinal analysis, spots from all ten D0–D3 captures were analyzed jointly. Harmony was applied to the principal components with capture identity as the batch variable ^35^, and the corrected representation was used to construct the neighborhood graph, UMAP embedding and Leiden clusters. The six expression clusters were displayed across time points and mapped back to the native coordinates of each capture. Cluster identity was evaluated using marker expression and the spatial distribution of spots.

For anatomical annotation of brain-section and replica maps, cluster locations were interpreted together with established regional markers and available tissue morphology. Selected gene-expression patterns were compared with ABA ISH images to identify corresponding brain structures.

### Spatial fidelity and transcriptomic concordance

Spatial expression profiles of *Neurod6*, *Hpca* and *Fibcd1* were used to assess signal spread near hippocampal boundaries by LWHM analysis ^20, 36^. Manually selected control points defined the high-expression contour, and spot distances were measured along its outward normal. Raw counts were averaged in distance bins, smoothed with a Gaussian filter and interpolated to obtain continuous profiles. Distances were converted using the 100-μm center-to-center pitch. LWHM was measured from the profile peak to the crossing at the midpoint between the peak and baseline on the evaluated side; profiles without a valid crossing were excluded.

For global transcriptomic concordance, gene counts were summed within each capture, matched by shared gene identifiers and transformed as log2(total counts + 1), followed by Pearson correlation. Gene-detection probability was the fraction of spots with a nonzero raw count, and logistic regression summarized its relationship with mean gene abundance. Transcript-length analyses used mean linear expression normalized to 10,000 counts per spot. After excluding low-expression genes and invalid length annotations, log2(Replica/Section) was related to log-transformed transcript length using locally weighted scatterplot smoothing (LOWESS).

For comparisons at matched sequencing depths, paired FASTQ reads were randomly subsampled with seqtk using a fixed seed to preserve pairing and then processed through the same alignment and counting workflow. Median UMI detection per tissue-covered spot, the proportion of valid barcodes, genomic read distribution and sequencing saturation were compared between transfer and direct-mounted preparations.

### Differential expression and functional enrichment

For Section–Replica comparisons, TBI and sham spots were compared separately within each preparation using the Wilcoxon rank-sum test in Scanpy on log-normalized expression. P values were adjusted by the Benjamini–Hochberg (BH) procedure ^37^, with adjusted P < 0.05 and |log2 fold change| > 1 defining differential expression. Mitochondrial, ribosomal, hemoglobin-associated, numerically designated Gm, pseudogene and Rik-labeled genes were excluded from the final gene lists. The 20 leading markers of TBI-section cluster 3 were evaluated in TBI-replica using the corresponding regressed and scaled expression values. Fold-change concordance between preparations was assessed using Pearson correlation across shared genes.

For longitudinal comparisons, raw counts were aggregated within each capture. Low-expression filtering with filterByExpr, trimmed mean of M-values (TMM) normalization ^38^ and limma-voom ^39^modeling were used for D1 versus D0, D2 versus D1, D3 versus D2 and D3 versus D0 comparisons. Differential-expression thresholds were BH-adjusted P < 0.05 and |log2 fold change| > 1. These analyses used captures from one animal as the observational units.

GO enrichment was performed separately for each analysis. Section–Replica comparisons used Enrichr for biological process, cellular component and molecular function terms ^40^. Longitudinal differential-expression and spatial-autocorrelation gene sets were evaluated using hypergeometric enrichment with BH correction and the corresponding tested genes as the background. The five temporal patterns were analyzed with g:Profiler using the genes eligible for temporal testing as the background and g:SCS correction ^41^. Coexpression-module enrichment used the network gene background, with BH correction over all tested module–term pairs. Terms with adjusted P < 0.05 were considered enriched; removal of redundant terms was performed only for presentation.

### Longitudinal gene expression analysis

Sample-level PCA used gene counts aggregated within each capture, converted to counts per million (CPM) and log2(CPM + 1)-transformed. Low-detection, mitochondrial and ribosomal genes were excluded, and the most variable genes were centered for PCA. To identify temporal patterns, log-normalized expression was averaged across all spots within each capture, including zero-expression spots. Genes meeting the expression filter in at least two captures were tested across D0–D3 by one-way analysis of variance (ANOVA) with BH correction, retaining 3,339 genes at adjusted P < 0.05.

Capture means were averaged with equal weights within each time point and Z-standardized across the four time points for each gene. Euclidean distance and Ward hierarchical clustering defined five temporal patterns. Trajectories show the mean standardized expression of member genes, with the inter-gene interquartile range as shading. Spatial cluster fractions were calculated within each capture and averaged with equal capture weights. Pattern scores within each cluster were adjusted for the capture-wide background, averaged across captures and standardized across clusters.

### Cell type deconvolution

Cell-type mapping of the brain replica was performed with cell2location using the mouse whole-brain single-nucleus reference dataset ^42^. Uncertain and low-quality reference labels were excluded, and cell-type expression signatures were estimated with reference sample identity included as a batch variable. Shared genes between the reference and spatial raw-count matrix were used for spatial mapping. The lower posterior quantile of inferred abundance was used for visualization of representative neuronal and glial populations.

For injury-associated analyses, RCTD, implemented in spacexr in full mode ^43^, was applied to SCC and D2-1 using a published single-cell reference of cortical SWI ^23^. Per-spot weights were normalized across all reference populations and summarized as relative RNA contributions. Reference-cell expression was normalized using each cell’s full-library UMI total and summarized by mean expression and detection fraction.

### Injury region mapping and C-SIDE analysis

SCC expression regions were annotated using the registered H&E image. Seurat SCTransform normalization ^44^ and anchor-based mapping ^45^ were used to transfer region labels from SCC to D2-1. Spots with a maximum prediction score of at least 0.5 received the predicted label; the remaining spots were marked unassigned. Euclidean distance from each D2-1 spot to the nearest C1-like spot center was measured in units of grid spacing and used to summarize regional RNA contributions.

Cell-type-specific expression associated with injury proximity was evaluated using C-SIDE while accounting for the RCTD-estimated composition ^46^. Proximity was defined as exp(−d/5), where d was the distance to C1 in grid spacings. BH correction was performed over all tested genes within each cell class, with a false discovery rate (FDR) < 0.01 and an absolute natural-log effect of at least 0.4 defining significant effects. Effects were converted to log2 units for display. Fitted expression was averaged within distance intervals and scaled to 0–1 by gene for the heatmap, retaining unavailable predictions as missing values.

### Spatial autocorrelation analysis

Moran’s I was calculated from log-normalized gene expression within each capture using row-normalized spatial weights linking directly adjacent grid positions. Low-detection genes and genes with insufficient valid autocorrelation estimates were excluded. Mean linear normalized expression and mean Moran’s I were calculated with equal capture weights at each time point, and D3–D0 changes were compared.

Two expression patterns were evaluated among genes with increased spatial autocorrelation: increased mean expression (log2 fold change ≥ 1) and relatively stable mean expression (|log2 fold change| < 0.25). To focus on the largest spatial changes, the D3–D0 increase in Moran’s I was ranked across all retained genes, and the top 10% were selected. Within this subset, 938 genes met the increased-expression criterion and 136 met the stable-expression criterion. These two gene sets were used for spatial comparisons and functional enrichment.

### Coexpression network construction and module activity analysis

Weighted gene coexpression analysis was performed on 13,795 genes retained after expression and technical-gene filtering. Spatially adjacent spots were aggregated into metaspots within each capture to reduce sparsity, and each capture contributed equal total weight to the analysis. Gene correlations were transformed into a signed adjacency matrix and then a topological overlap matrix (TOM) ^47^. Hierarchical clustering based on TOM dissimilarity, followed by dynamic tree cutting ^48^ and merging of related modules, identified six coexpression modules, M01–M06.

Module activity at individual spots was calculated by projecting expression onto module-specific first-principal-component loadings derived from metaspots. Scores were standardized with equal total weight per capture and summarized by time point and expression region; spatial organization was assessed using within-region Moran’s I. Hub genes were ranked by their correlation with the module eigengene, and the 15 leading M02 hubs were visualized using their pairwise TOM similarities.

### Statistical analysis and reproducibility

Statistical tests were selected for each comparison as specified above and in the figure legends. PI-positive capture area and total RNA recovery were measured in three repeated preparations from the same tissue for each condition and summarized as mean ± standard error of the mean (s.e.m.). Group comparisons used one-way ANOVA with Tukey’s multiple-comparisons test. Transcriptomic concordance was quantified using Pearson correlation, and multiple gene-level testing used the correction procedures and thresholds specified for each analysis. Analysis and visualization were implemented in Python and R.

## Author Contribution

X.Z. and Z.Y. conceived and supervised the study. W.W. developed the nanoforest interface, performed the sampling experiments and wrote the first draft. K.Y. fabricated and characterized the barcoded capture chips. X.M., K.D. and Y.H. performed library preparation and sequencing. H.W. and L.H. performed the animal surgery and histological analysis. F.D. provided neurosurgical expertise and contributed to the injury model. L.C. and H.L. contributed to clinical interpretation and sample strategy. All authors analyzed data, discussed the results and commented on the manuscript.

## Acknowledgements

This work was financially supported by National Natural Science Foundation of China grant 82361138570 (X.Z.), 81827901 (X.Z.), 32602071 (K.Y.), Southeast University Innovation Capability Enhancement Plan for Doctoral Students grant CXJH_SEU24139 (W.W.), and Postgraduate Research &Practice Innovation Program of Jiangsu Province KYCX23_0312 (W.W.).

## Data and Code Availability

Raw sequencing data generated in this study have been deposited in the Genome Sequence Archive (GSA) of the National Genomics Data Center under accession number CRA087453. Processed spatial transcriptomic data, including gene-expression matrices, spatial-spot metadata, and spatial coordinates in AnnData format, are publicly available through Zenodo (https://doi.org/10.5281/zenodo.24160470). The mouse whole-brain single-nucleus RNA-sequencing dataset used for cell-type deconvolution is publicly available from ArrayExpress under accession number E-MTAB-11115. The mouse cerebral cortex single-cell RNA-sequencing dataset used for injury-related analyses is publicly available from the Gene Expression Omnibus (GEO) under accession number GSE226207. Allen Brain Atlas in situ hybridization (ISH) data used for spatial validation are publicly available from the Allen Brain Atlas. Custom FASTQ preprocessing scripts and key analysis code used in this study have been permanently archived in Zenodo at https://doi.org/10.5281/zenodo.24187342. The actively maintained version is additionally available at https://github.com/Wong-WJ/LIFE-seq. Custom microneedle arrays and nanoforest-based capture interfaces were fabricated as described in the Materials and Methods, and detailed information required for their reproduction is provided in the manuscript and Supplementary Materials. All other data needed to evaluate and reproduce the results presented in this study are provided in the paper and/or the Supplementary Materials.

