## Supplementary information for "LIFE-seq: longitudinal spatial transcriptomics of living tissue"

### Extended Data

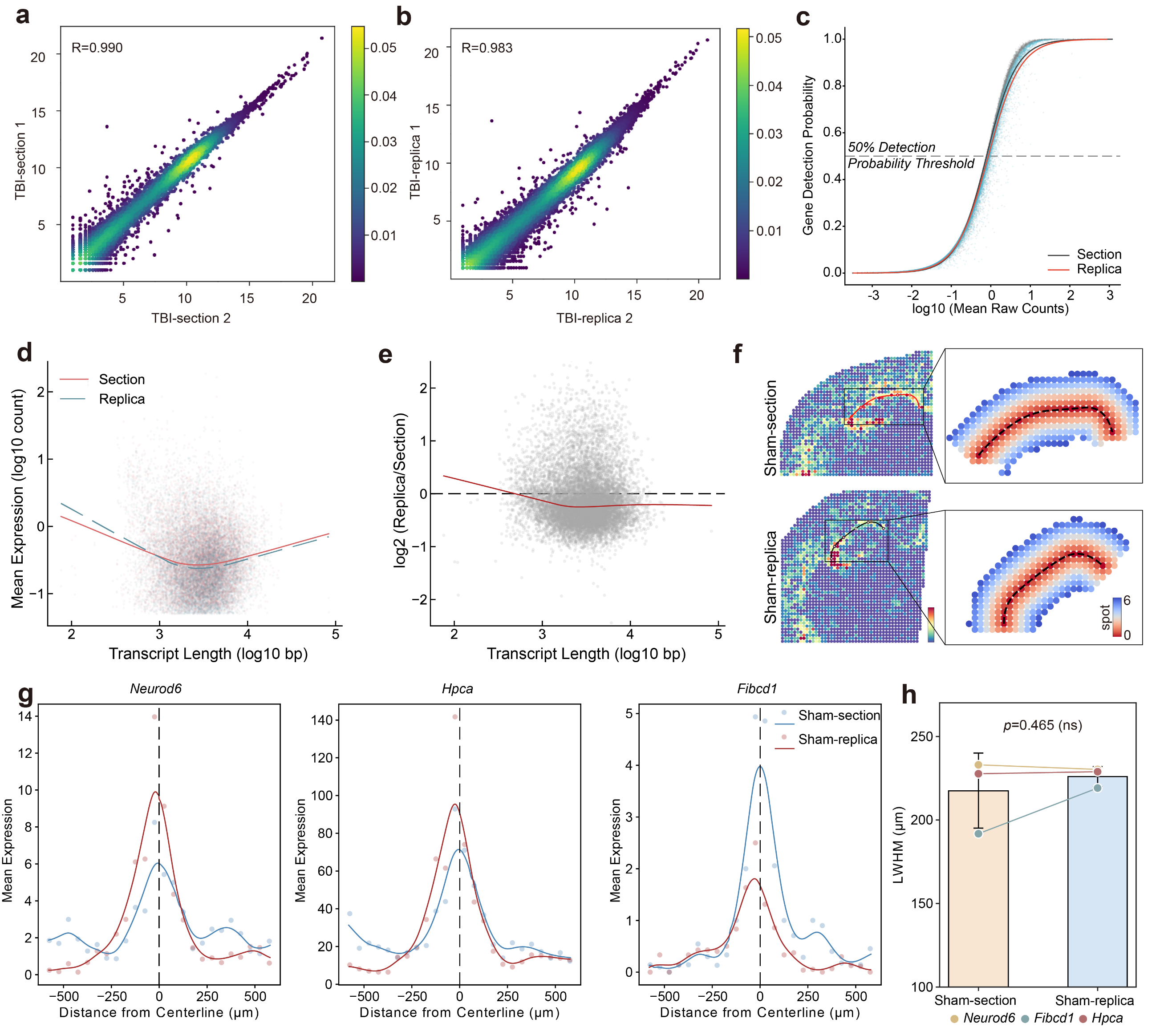

**Extended Data Fig. 1** **Quality control and spatial resolution validation of LIFE-seq in the TBI model. a, b,** Scatter plots illustrating the extremely high transcriptomic reproducibility between two independent replicates for the TBI-section (**a,** R = 0.99) and TBI-replica (**b,** R = 0.983) groups. **c,** Scatter plot evaluating the gene detection probability (fraction of spots) as a function of expression abundance (log10 mean raw counts) for the Section and Replica groups. **d, e,** Evaluation of transcript length bias. Mean expression (log10 count) (**d**) and the log2-transformed expression ratio of Replica to Section (**e**) plotted as a function of transcript length (log10 bp). **f,** Spatial visualization of the region-restricted gene *Hpca* and the corresponding fitted tissue boundary lines in Sham-section and Sham-replica, showing the selected baseline and the normal vector traversing 6 spots outward. **g,** LWHM diffusion curves for three representative genes (*Neurod6*, *Hpca*, *Fibcd1*) in the Sham-section and Sham-replica groups. **h,** Bar plot showing the left-width at half-maximum (LWHM) of expression intensity around the boundary line for *Neurod6*, *Hpca*, and *Fibcd1*. Data are represented as mean ± 95% confidence interval (CI). The P value was calculated using a pairwise t-test.

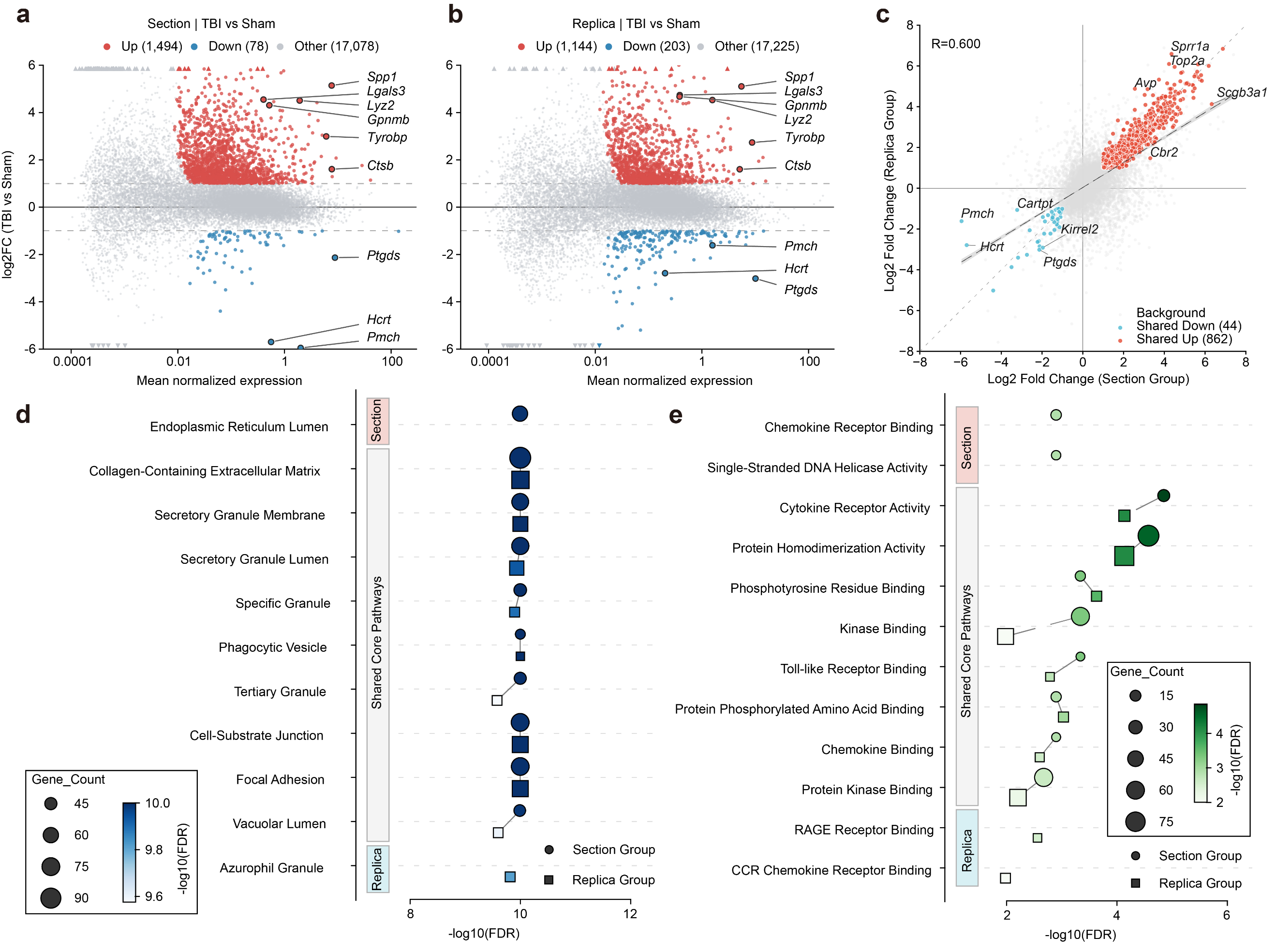

**Extended Data Fig. 2 Transcriptomic profiling and pathway enrichment of traumatic brain injury using LIFE-seq.** **a, b,** MA plots showing mean normalized gene expression (x-axis) and log2 fold changes (y-axis) for TBI versus Sham comparisons in the Section (**a**) and Replica (**b**) groups. Red, blue and grey indicate upregulated, downregulated and other genes, respectively. **c,** Scatter plot showing the correlation of log2 fold changes between the Section (x-axis) and Replica (y-axis) groups (R = 0.6), highlighting 862 shared upregulated and 44 shared downregulated genes. **d, e,** Comparison of the top 10 enriched Gene Ontology (GO) Cellular Component (CC, **d**) and Molecular Function (MF, **e**) terms for upregulated DEGs in the Section and Replica groups. Circles and squares represent the Section and Replica groups, respectively. Bubble size indicates gene count, and color represents −log10(FDR). Nine overlapping CC terms and eight overlapping MF terms are shown.

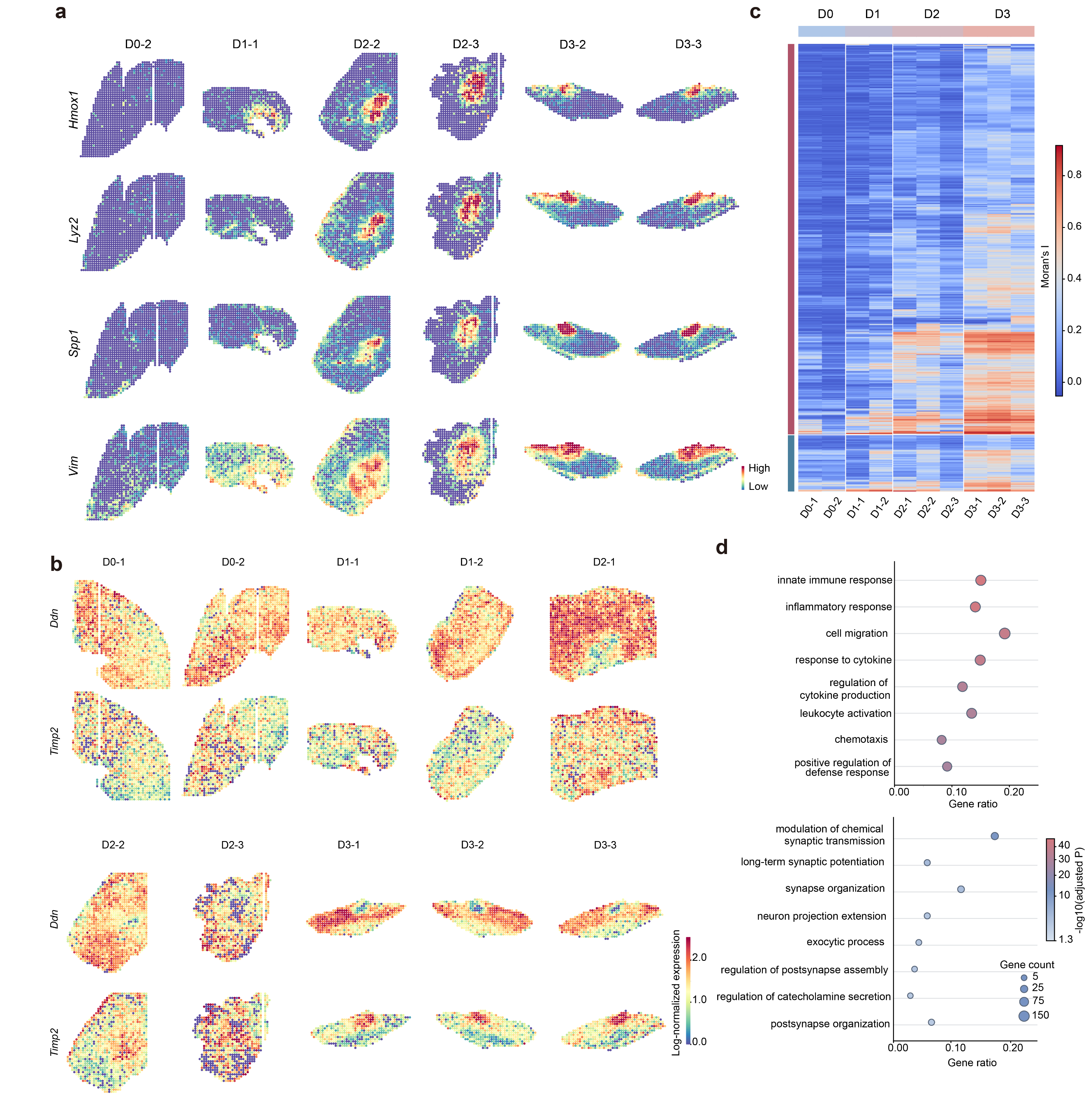

**Extended Data Fig. 3 Spatiotemporal expression patterns and spatial autocorrelation of injury-associated genes.** **a,** Spatial expression maps of four representative injury-responsive genes (*Hmox1*, *Lyz2*, *Spp1* and *Vim*) across six samples from D0–D3. **b,** Spatial expression maps of *Ddn* and *Timp2* across all ten longitudinal samples. Colors indicate log-normalized expression levels. **c,** Heatmap showing Moran's I values for representative genes across ten samples, grouped into genes with increased expression and spatial autocorrelation (top) and genes with relatively unchanged expression but increased spatial autocorrelation (bottom). **d,** Gene Ontology (GO) Biological Process (BP) enrichment analysis of the two gene groups, showing immune- and inflammation-related processes (top) and neuronal and synaptic processes (bottom). Bubble size indicates gene count, color represents −log10-adjusted P value, and the x-axis indicates gene ratio.

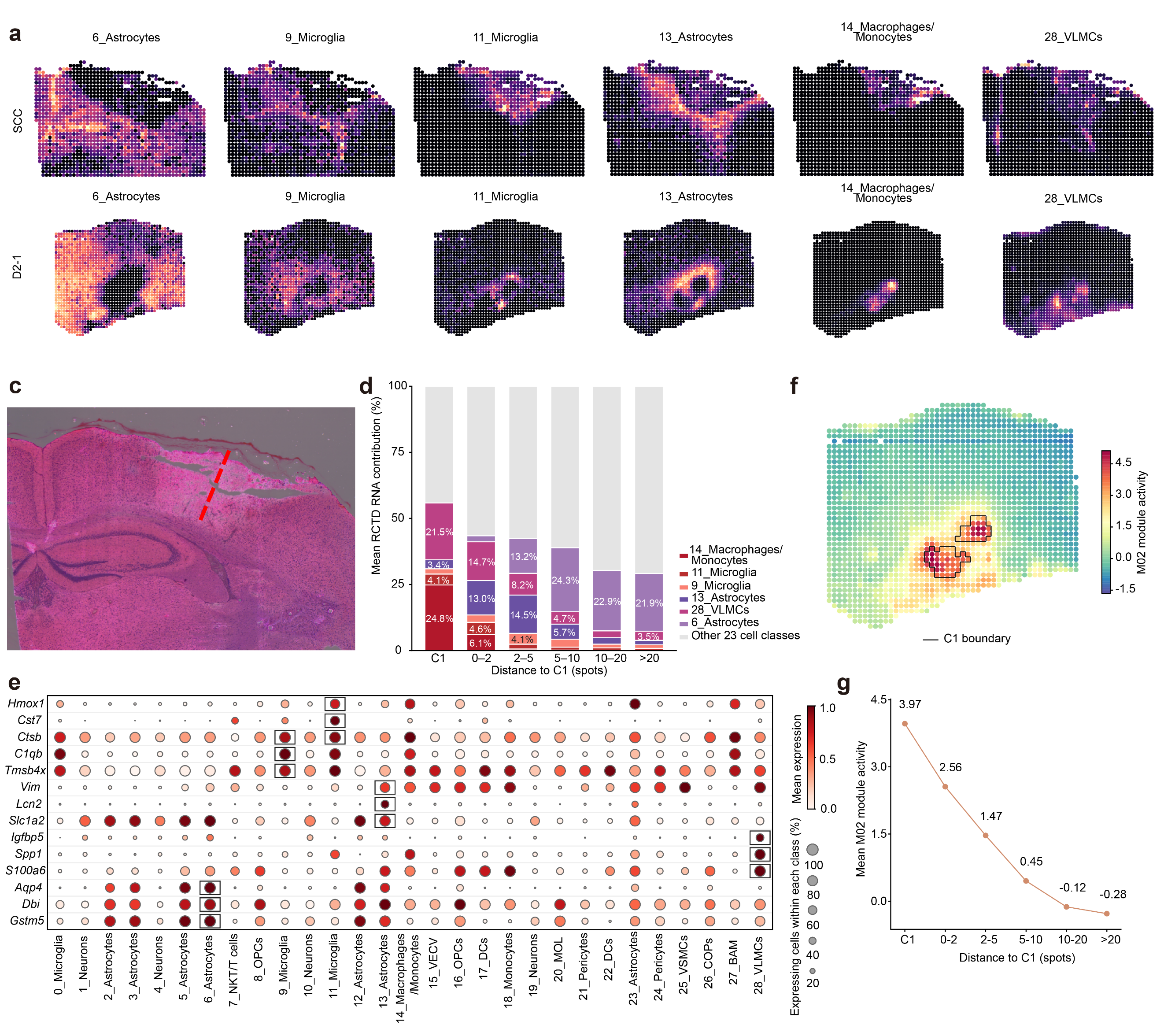

**Extended Data Fig. 4 Cell-type composition and spatial distribution of injury-associated gene programs.** **a, b,** Spatial maps of RCTD-estimated RNA contributions for six representative cell populations in the endpoint coronal section (SCC, **a**) and D2-1 (**b**), including astrocytes, microglia, macrophages/monocytes and vascular and leptomeningeal cells (VLMCs). **c,** Hematoxylin and eosin (H&E) staining of the endpoint coronal section, with the red dashed line indicating the injury site. **d,** Stacked bar plots showing mean RCTD-estimated RNA contributions of major cell types within C1 and surrounding regions stratified by distance from C1. **e,** Dot plot showing expression of representative marker genes across 29 cell populations in the single-cell reference. Dot size indicates the percentage of expressing cells, and color represents mean expression. Boxes highlight selected cell-type-specific markers. **f,** Spatial distribution of M02 module activity in D2-1, with the C1 boundary outlined in black. **g,** Mean M02 module activity across C1 and five surrounding distance intervals (0–2, 2–5, 5–10, 10–20 and >20 spots).

### Supplementary Data

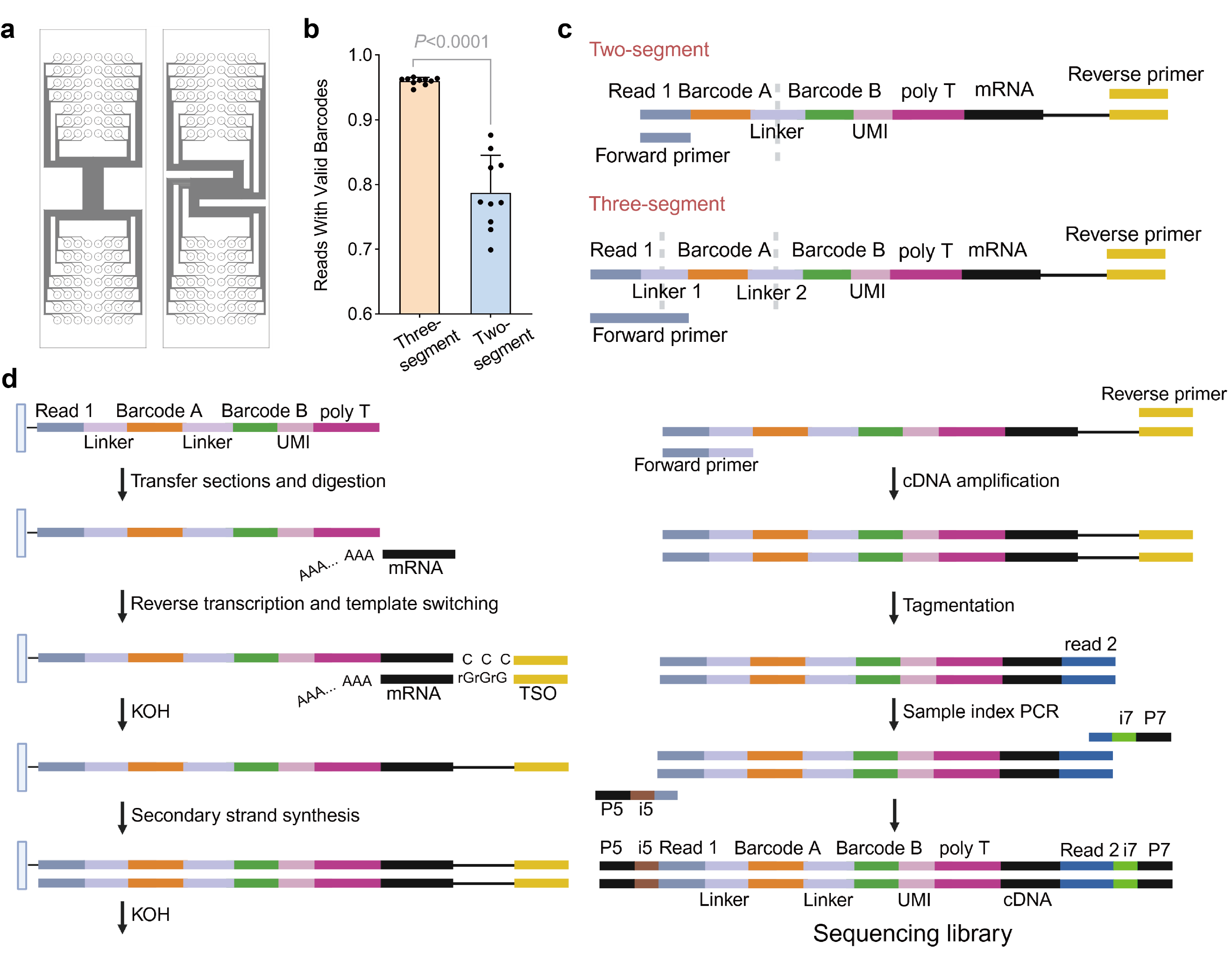

**Supplementary Fig. S1 Design of microfluidic orthogonal barcoding chips and workflow of three-segment probe assembly and library construction.** **a,** Layout design of a pair of polydimethylsiloxane (PDMS) microfluidic channel chips used for sequential orthogonal spatial barcoding. **b,** Comparison of valid spatial barcode detection rates between the three-segment and two-segment probe assembly strategies (*P* < 0.0001). **c,** Schematic illustration of the probe architectures, ligation junctions (indicated by white dashed lines), and primer-binding sites for the two-segment and three-segment barcoding strategies. **d,** Schematic workflow of LIFE-seq library construction, sequentially depicting tissue transfer and *in situ* digestion, mRNA poly(A) tail capture, *in situ reverse transcription and template switching*, KOH denaturation, second-strand cDNA synthesis, cDNA pre-amplification, Tn5 tagmentation, and sample index PCR.

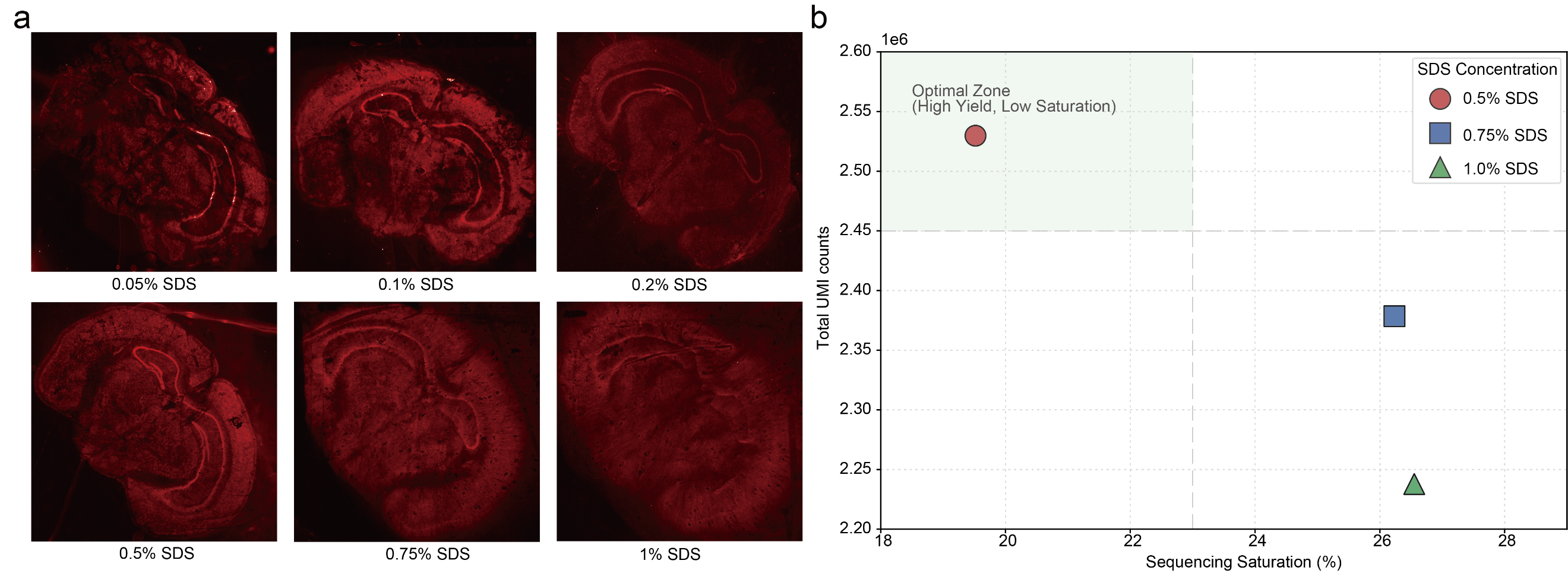

**Supplementary Fig. S2 Optimization of SDS concentration in the in situ tissue digestion buffer.** a, Fluorescence microscopy images of cDNA footprints on the barcoding chip obtained at different SDS concentrations. b, Total UMI counts plotted against sequencing saturation at SDS concentrations of 0.5%, 0.75% and 1.0%. The shaded region highlights conditions with high total UMI counts and low sequencing saturation.

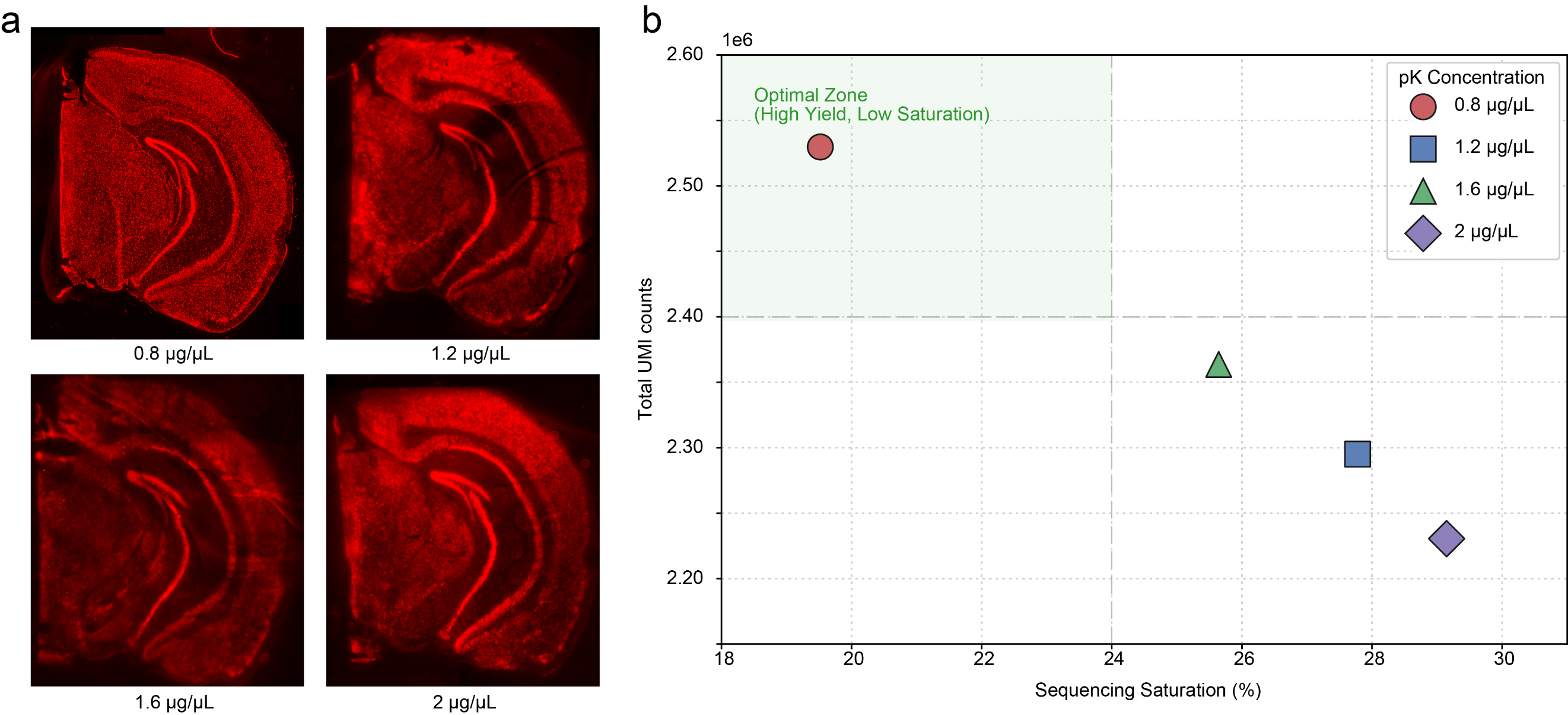

**Supplementary Fig. S3 Optimization of proteinase K (PKD) concentration in the in situ tissue digestion buffer.** a, Fluorescence microscopy images of cDNA footprints on the barcoding chip obtained at different proteinase K concentrations. b, Total UMI counts plotted against sequencing saturation at proteinase K concentrations of 0.8, 1.2, 1.6 and 2.0 μg/μL. The shaded region highlights conditions with high total UMI counts and low sequencing saturation.

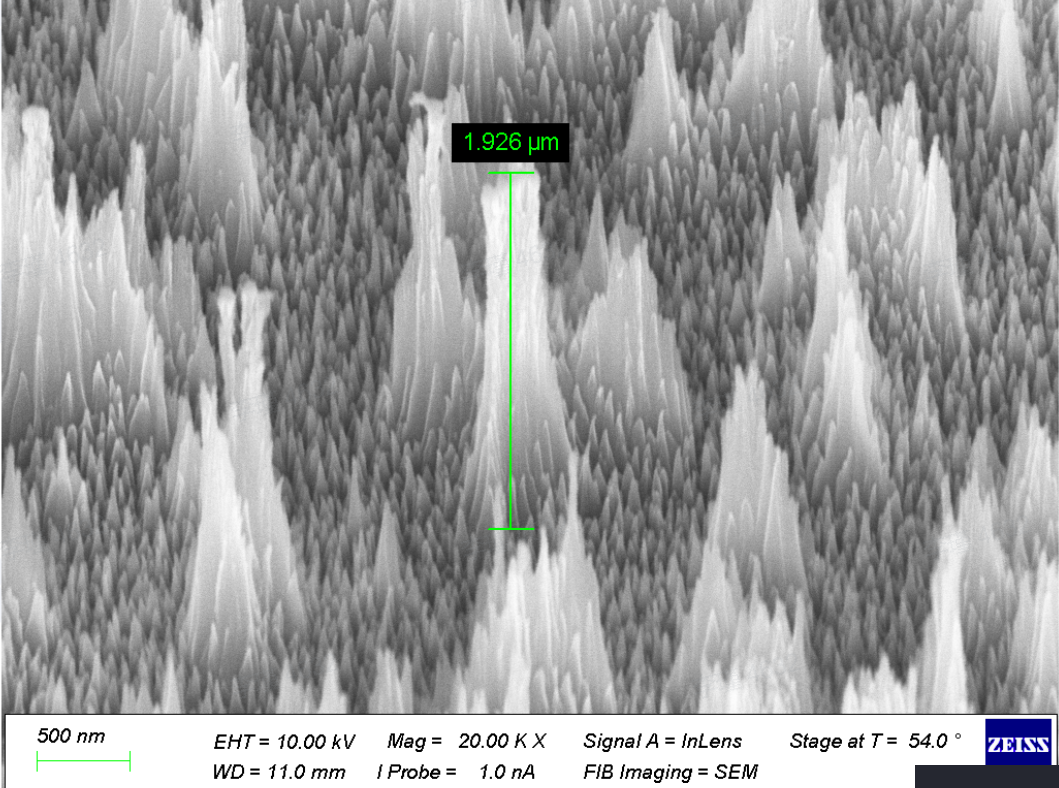

**Supplementary Fig. S4 Morphological characterization of the quartz nanoforest array.** Tilted scanning electron microscopy (SEM) image showing the densely packed, disordered quartz nanoforest array with high-aspect-ratio nanostructures. The measured height of the representative nanostructure is 1.926 μm, consistent with the nominal array height of approximately 2 μm. Scale bar, 500 nm.

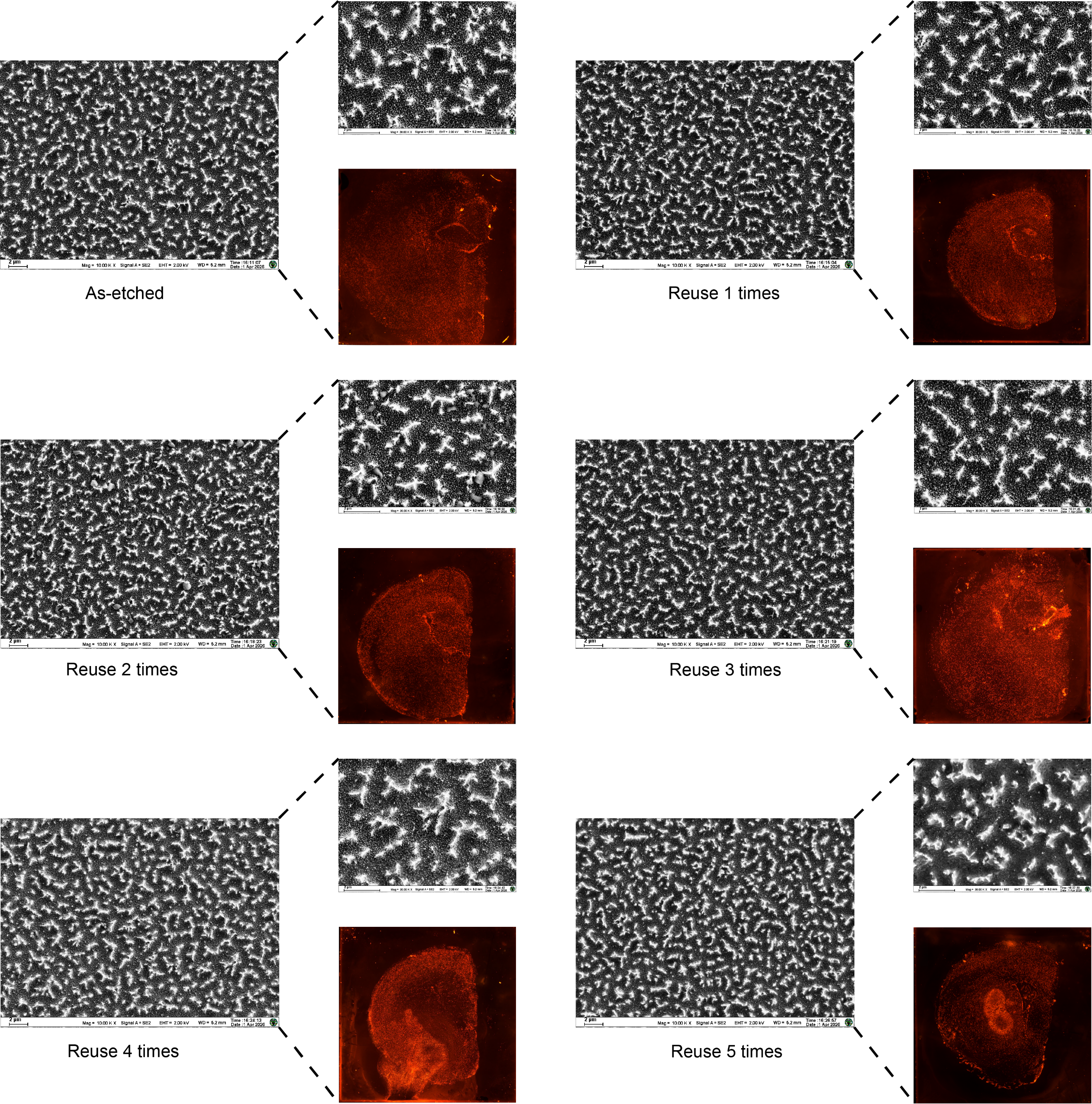

**Supplementary Fig. S5 Morphological stability and reproducibility of nanoforest-mediated tissue sampling.** Representative scanning electron microscopy (SEM) images of nanoforest arrays at low (left) and high (upper right) magnifications, together with corresponding propidium iodide (PI) fluorescence images of captured mouse brain tissue replicas (lower right) across six repeated sampling experiments. The SEM images show consistent nanoforest morphology, while the PI fluorescence images demonstrate reproducible tissue capture with preserved anatomical outlines. Scale bars are indicated in the SEM images.

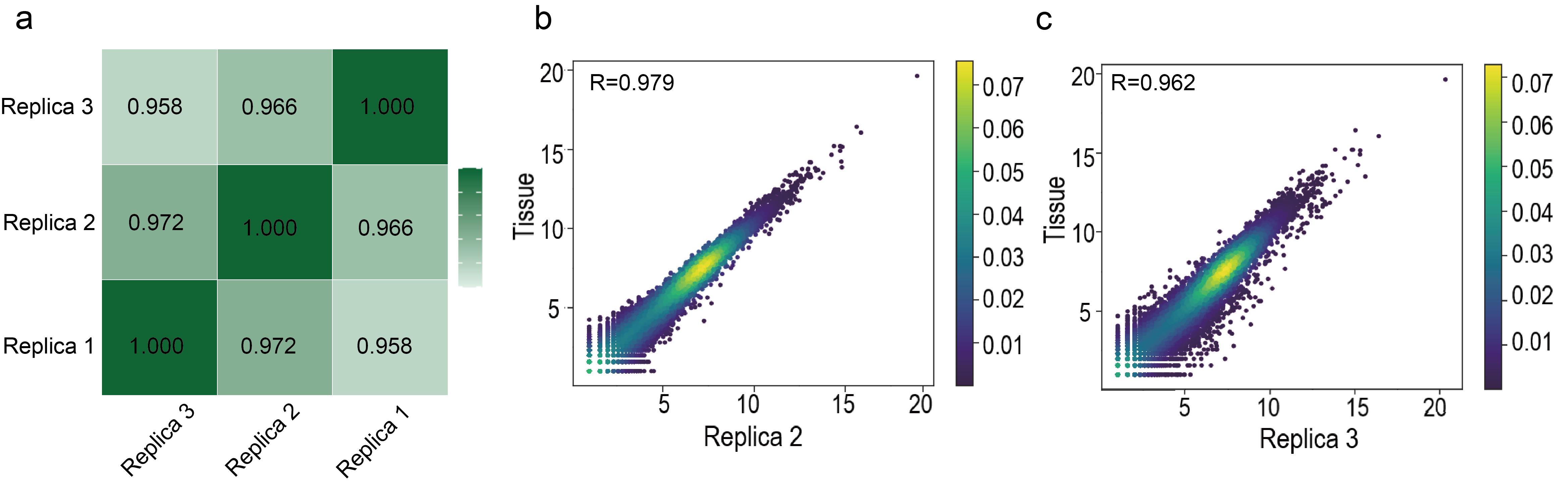

**Supplementary Fig. S6 Transcriptomic reproducibility of nanoforest-derived tissue replicas.** **a,** Pearson correlation matrix comparing global gene expression profiles among three nanoforest replicas (Replica 1–3), with pairwise correlation coefficients ranging from 0.958 to 0.972. **b, c,** Density scatter plots showing the correlations of log-transformed gene expression between the native tissue section (Tissue, y-axis) and Replica 2 (b, x-axis, R = 0.979) or Replica 3 (c, x-axis, R = 0.962). Point colors indicate local density.

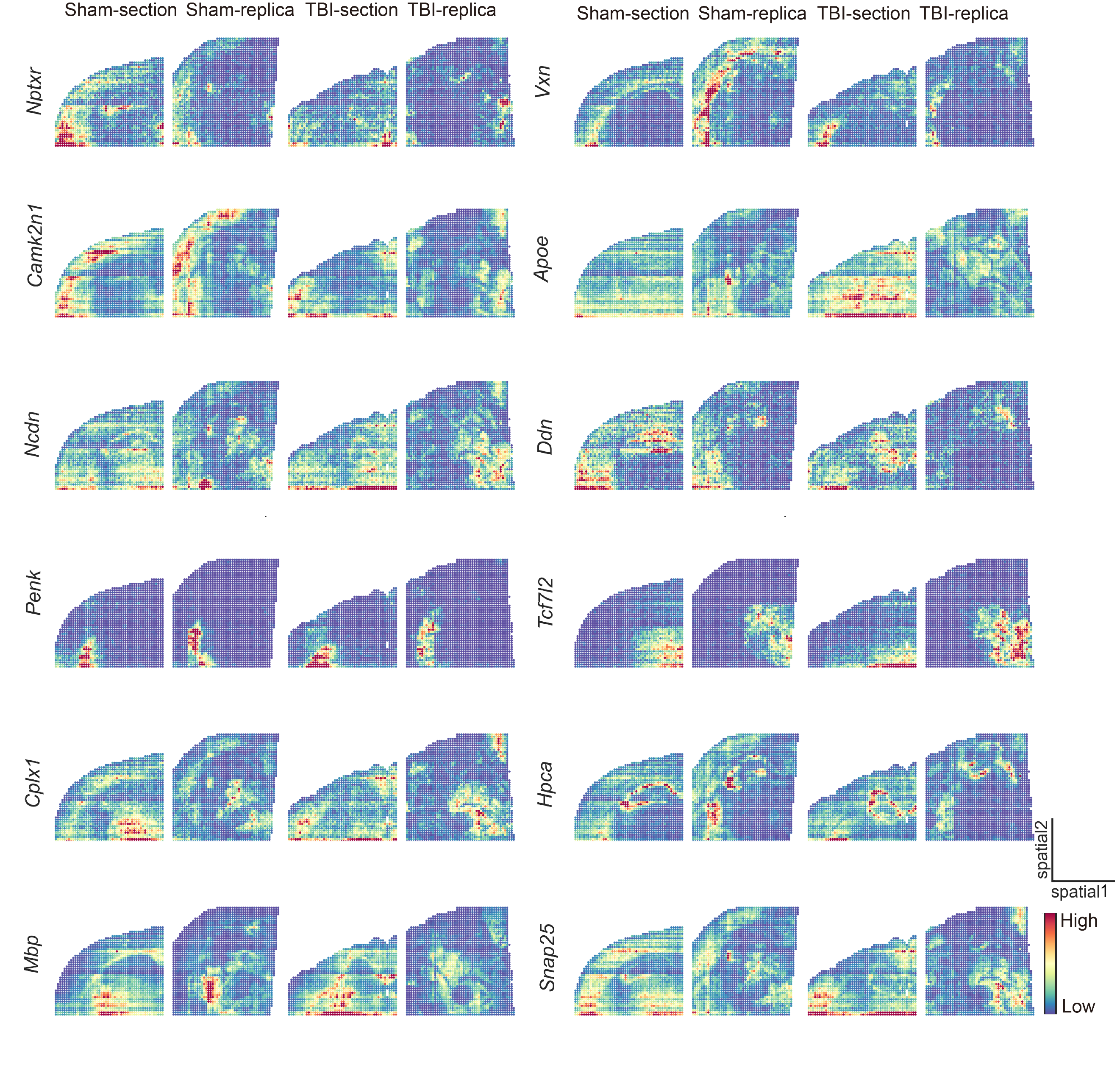

**Supplementary Fig. S7 Spatial expression patterns of representative anatomical marker genes in tissue sections and nanoforest replicas.** Spatial expression maps of 12 representative brain marker genes (*Nptxr*, *Camk2n1*, *Ncdn*, *Penk*, *Cplx1*, *Mbp*, *Vxn*, *Apoe*, *Ddn*, *Tcf7l2*, *Hpca* and *Snap25*) across four groups: Sham-section, Sham-replica, TBI-section and TBI-replica. Color intensity represents relative gene expression levels, ranging from low (purple) to high (yellow–red). Corresponding Section and Replica samples exhibit consistent anatomical expression patterns under both Sham and TBI conditions, supporting the spatial fidelity of nanoforest-mediated molecular sampling.

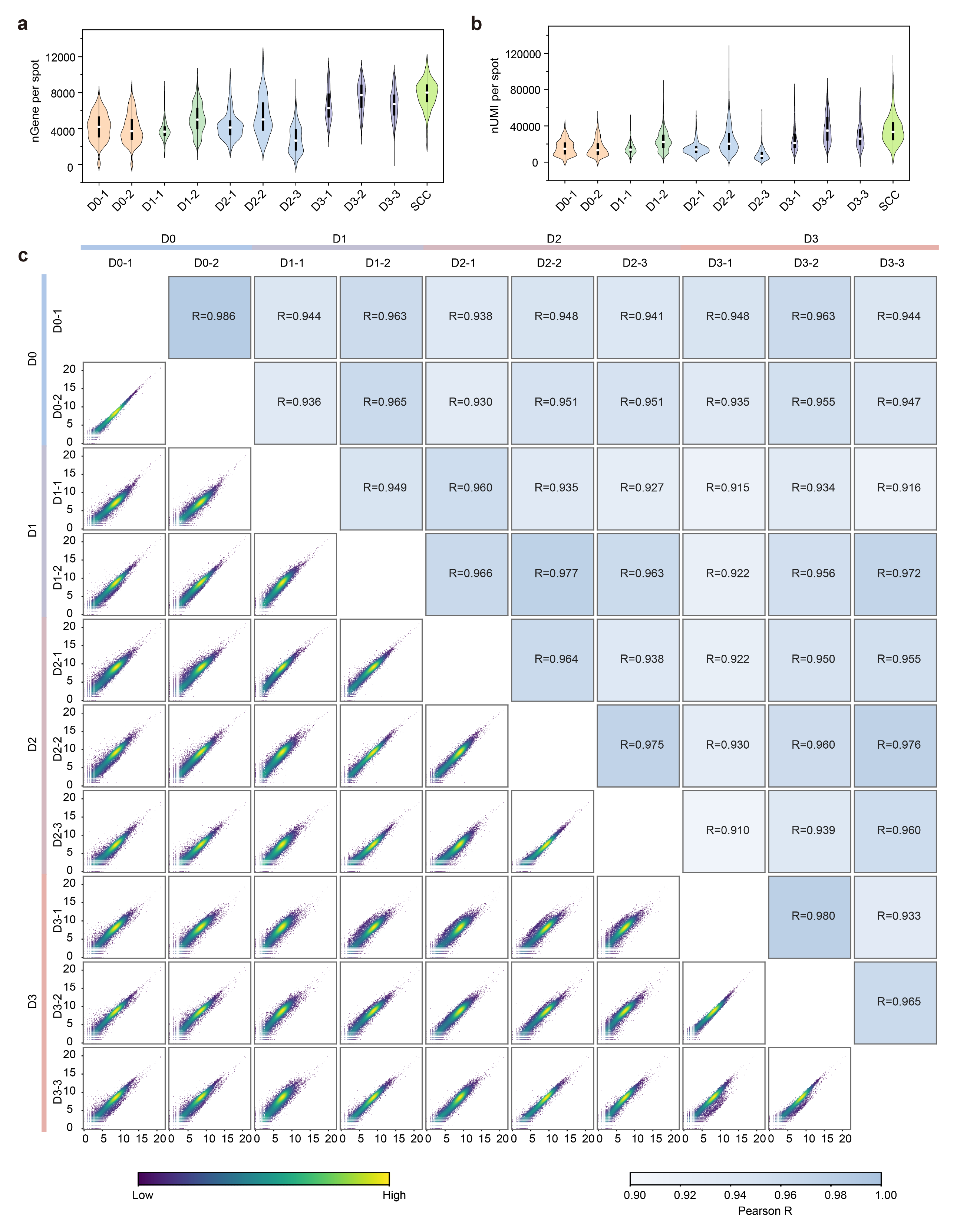

**Supplementary Fig. S8 Molecular detection and transcriptomic correlations in longitudinal cortical injury samples.** **a, b,** Violin plots showing the numbers of detected genes (**a**) and UMIs (**b**) per spot across ten longitudinal samples and the SWI coronal control (SCC). **c,** Pairwise Pearson correlations of global gene expression across ten samples. The upper triangle shows correlation coefficients, and the lower triangle shows gene-expression density scatter plots colored by local density. D0–D2 samples were collected in vivo using LIFE-seq, whereas D3 samples were obtained by conventional tissue-section mounting after euthanasia.

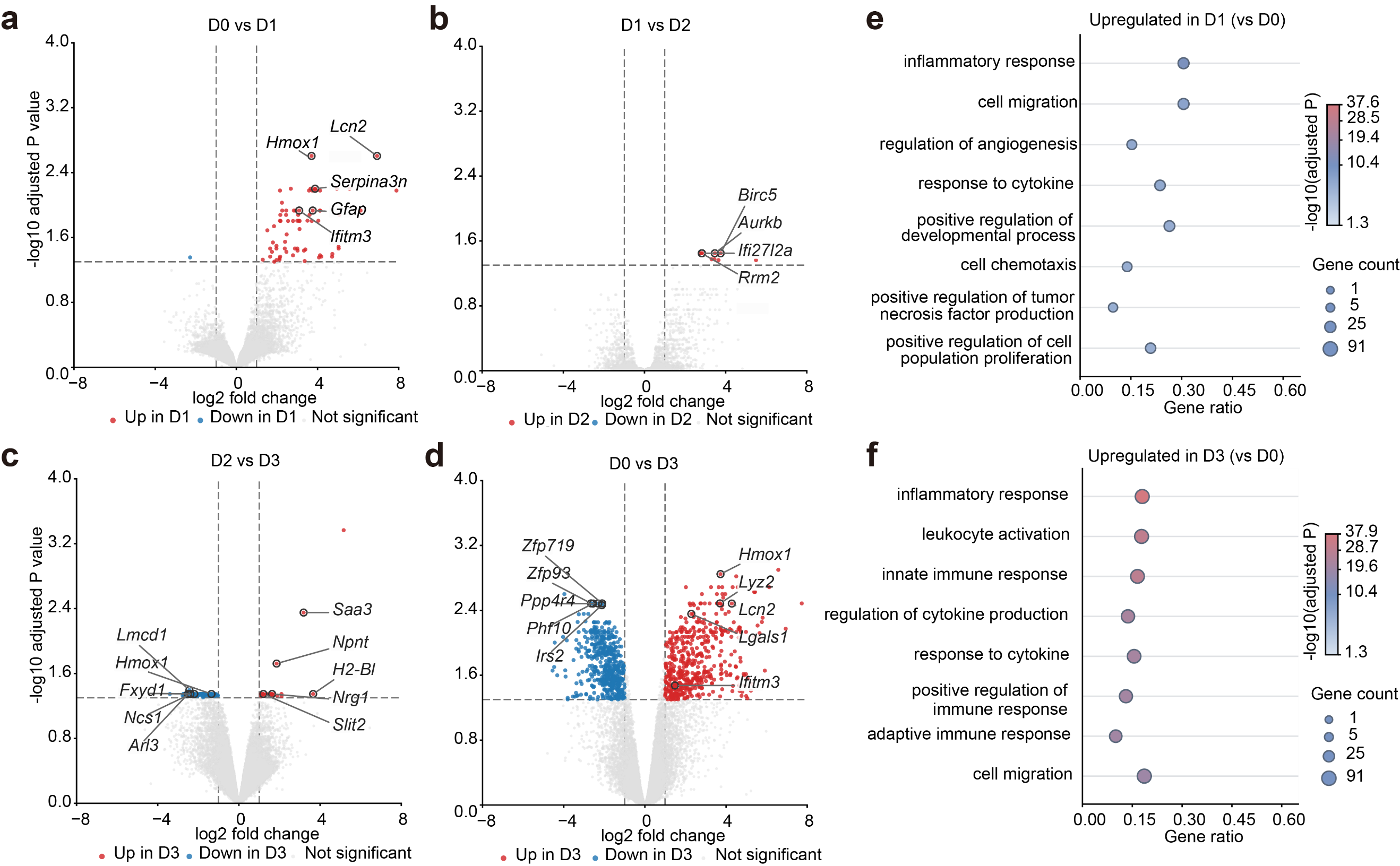

**Supplementary Fig. S9 Stage-associated expression changes and functional enrichment following cortical injury. a–d,** Volcano plots comparing D1 versus D0 (a), D2 versus D1 (b), D3 versus D2 (c) and D3 versus D0 (d). The x-axis indicates log2 fold change, and the y-axis indicates −log10-adjusted P value. Red, blue and grey denote upregulated, downregulated and non-significant genes, respectively; selected genes are labeled. Differentially expressed genes were defined by adjusted P<0.05 and |log2FC|>1. Analyses used raw counts aggregated within each sample, followed by TMM normalization and limma-voom modeling. **e,f,** Representative enriched GO biological processes among genes upregulated at D1 versus D0 (e) and D3 versus D0 (f). The x-axis indicates the fraction of effectively mapped input genes assigned to each term. Bubble size represents the number of associated genes, and color indicates −log10-adjusted P value. Enrichment was assessed using a hypergeometric test with the tested genes as background and Benjamini–Hochberg correction. D0–D2 samples were collected in vivo using LIFE-seq; D3 samples were obtained by endpoint tissue-section mounting after euthanasia.

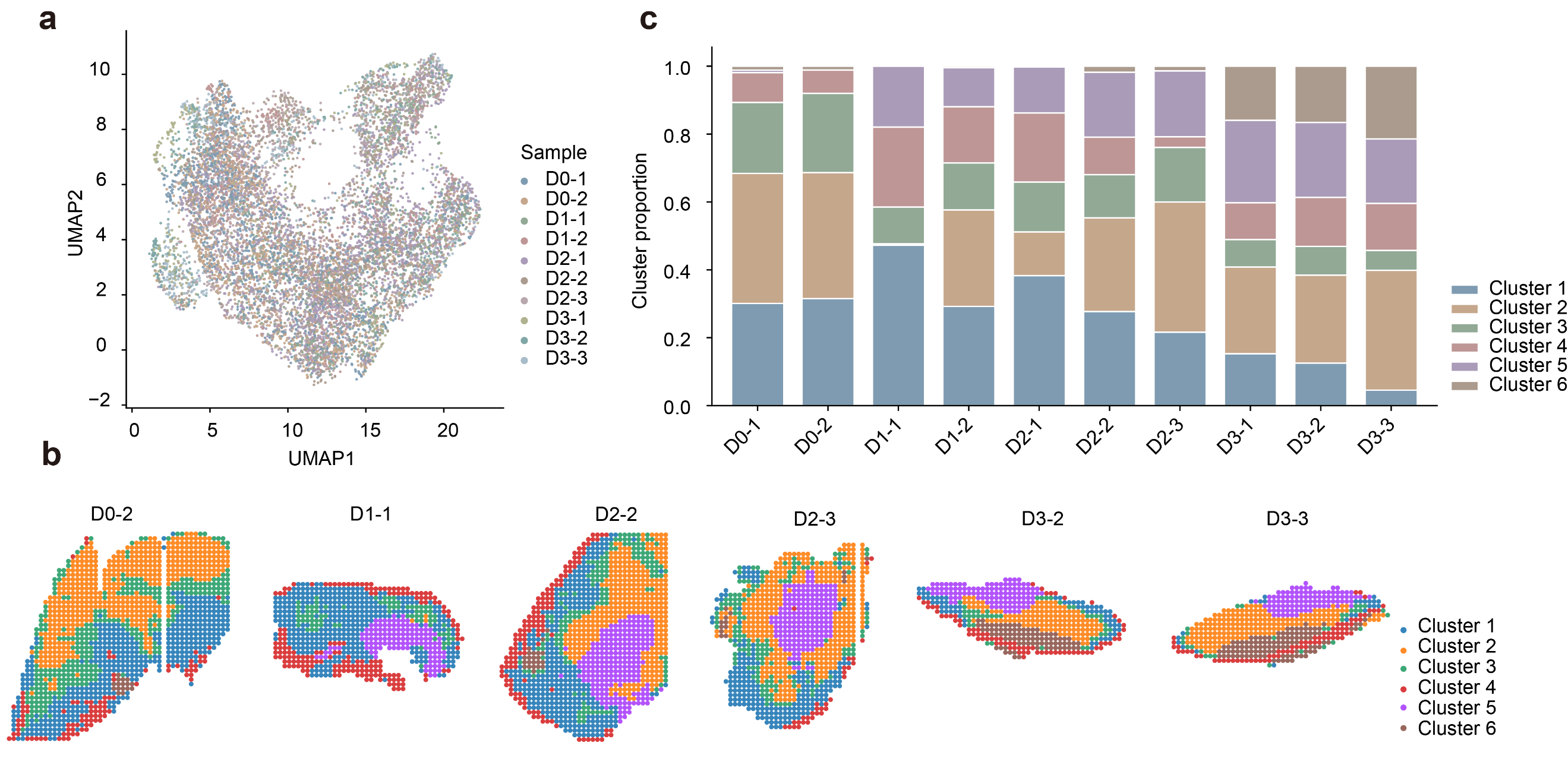

**Supplementary Fig. S10 Spatial clustering and cluster composition of longitudinal cortical injury samples.** **a,** Joint UMAP of all ten spatial transcriptomic samples, colored by sample identity. **b,** Spatial distributions of six expression clusters in the remaining six samples (D0-2, D1-1, D2-2, D2-3, D3-2 and D3-3). Colors indicate cluster assignments. **c,** Stacked bar plots showing the proportions of six expression clusters in each of the ten samples. All samples originated from the same mouse.

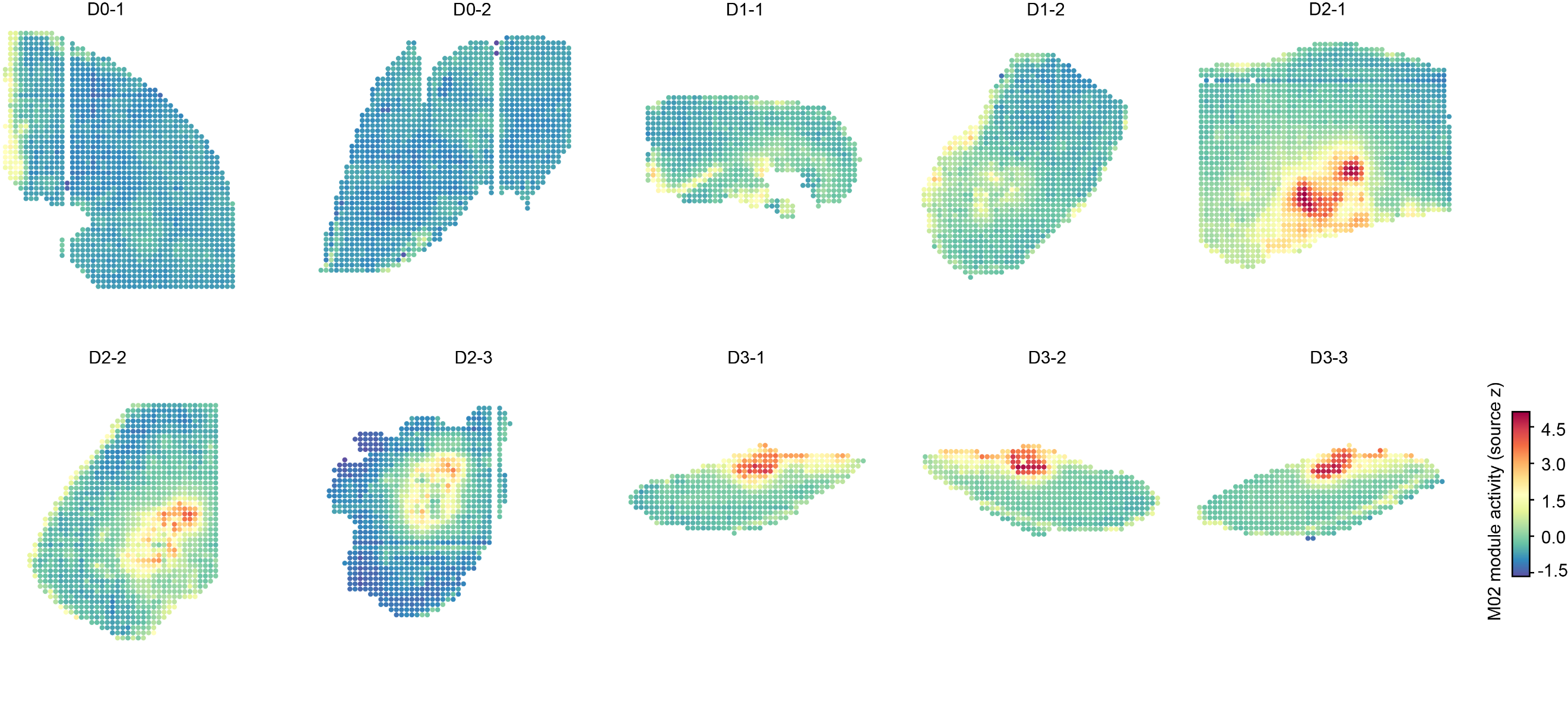

**Supplementary Fig. S11 Spatiotemporal distribution of M02 coexpression module activity following cortical injury.** Spatial maps of M02 module activity across ten samples from D0–D3. Colors indicate source-standardized module activity (Z-score).
